# SGLT2 Inhibition Ameliorates Age-Dependent Renovascular Rarefaction

**DOI:** 10.1101/2025.06.27.654312

**Authors:** Anastasia Paulmann, Matthew D. Cox, Tom Boewer, Hannah M. Somers, Heath Fuqua, Ryan P. Seaman, Joel H. Graber, Anchal Mahajan, Cory P. Johnson, Laura L. Beverly-Staggs, Sonia Sandhi, Heiko Schenk, Hermann Haller

## Abstract

**Introduction:** Aging is associated with progressive loss of renal function and vascular structure, with and without chronic kidney disease. However, the mechanisms driving renal vascular aging and potential therapeutic interventions remain poorly understood.

**Methods:** To model this state-of-affairs, we used African turquoise killifish (Nothobranchius furzeri), a naturally short-lived vertebrate. We then inhibited the sodium-glucose co-transporter 2 using dapagliflozin (SGLT2i) to test a potential therapeutic intervention. Histological, immunofluorescent, and 3D vascular imaging were used to evaluate glomerular, tubular, vascular and functional changes. Single-nuclei transcriptomic profiling was performed on whole kidneys to identify age- and treatment-associated molecular signatures.

**Results:** Aged killifish kidneys exhibited hallmark features of renal aging, including glomerulosclerosis, tubular fibrosis, and vascular rarefaction. Functional changes included increased proteinuria and altered tubular transporter function. Transcriptomic profiling revealed a metabolic shift from oxidative phosphorylation to glycolysis and upregulation of pro-inflammatory pathways. Aged vasculature also displayed a marked reduction in tight junctions and cell–cell contacts. SGLT2i attenuated age-related vascular rarefaction, preserved functional capillary networks, reduced albuminuria, restored a youthful transcriptional profile and enhanced intercellular signaling. However, killifish lifespan was not extended.

**Conclusion:** This study establishes the killifish as a translational model for investigating renal vascular aging. We show that SGLT2i preserves renal microvascular structure and function, reduces proteinuria, and reprograms the aged transcriptome. These results support a vascular-protective role of SGLT2i in mitigating age-related renal deterioration.

**Translational Statement:** This study establishes the African turquoise killifish as a model for investigating renal and vascular aging. We found that SGLT2 inhibition preserves microvascular integrity and reduces proteinuria. These results mirror established benefits observed in mammalian models and patients with chronic kidney disease, reinforcing the kidney-protective role of SGLT2 inhibitors. However, the killifish offers a unique opportunity for rapid, translational aging research. By using a naturally short-lived vertebrate with mammalian-like renal aging, our model enables a rapid, preclinical, assessment of vascular outcomes and identifies microvascular preservation as a potential mechanistic target for renoprotection.

## Introduction

The kidney is particularly susceptible to age-associated deterioration. In patients with chronic kidney disease (CKD), aging accelerates disease progression, impacting approximately 10% of adults worldwide ^1^. However, therapies addressing molecular mechanisms to prevent or delay kidney aging are limited.

Renal aging features a loss of functional nephrons, tubulointerstitial fibrosis, and microvascular rarefaction ^2^. These changes collectively increase the susceptibility of patients to other age-associated conditions including hypertension and metabolic disease. Yet the mechanisms underlying these renal changes remain incompletely understood. Microvascular rarefaction, a reduction in the density of glomerular and peritubular capillaries, is increasingly recognized as a pivotal contributor to age-related renal decline ^3^. Capillary loss promotes local hypoxia, disrupts metabolic homeostasis, and exacerbates fibrosis, creating a self-reinforcing loop of progressive damage. Therapeutically targeting the renal microvasculature presents a promising approach to preserving kidney health with age.

Furthermore, modeling aging longitudinally is challenging, time- and cost-intensive. Nonvertebrate models lack key mammalian-like organ systems, including vasculature and kidneys. Traditional vertebrate models such as mice and have relatively long lifespans (2–5 years), limiting their utility ^4^. The African turquoise killifish (*Nothobranchius furzeri*; ATK) has emerged as a powerful vertebrate model organism due to its naturally short lifespan of just 4–6 months and its vertebrate-typical organ systems ^5^; including a kidney structurally comparable to that of humans. Despite increasing use in aging research, the aging killifish renal system has not been systematically characterized, and its vasculature has not been examined.

Sodium-glucose linked transporter 2 inhibitors (SGLT2i) have emerged as remarkable agents improving renal outcomes in both diabetic and non-diabetic patients. Clinical studies have shown that SGLT2i improves cardiac and renal function ^6^, support brain health ^7^, while also significantly enhances survival in patients with chronic diseases^8^, many of which are associated with microvascular deterioration ^9,10^. Despite SGLT2i’s growing relevance, little is known about their potential anti-aging effects or the underlying molecular mechanisms.

We hypothesized that killifish could facilitate insights into the aging kidney and that SGLT2i could provide a model therapeutic intervention. We used conventional methods coupled with single-nucleus RNA sequencing to resolve cell type-specific transcriptome changes during aging and applied CellChat^11^ to explore alterations in intercellular signaling. We present the first systematic characterization of renal aging in ATK, focusing on vascular rarefaction and age-related proteinuria onset. We found that aging is associated with progressive microvascular rarefaction and glomerular dysfunction, and that SGLT2i treatment preserves vascular integrity and is associated with a lower incidence of proteinuria in aged fish.

## Methods

Detailed protocols are available in the supplementary materials. This manuscript was prepared in accordance with the ARRIVE 2.0 guidelines^12^, and the ARRIVE reporting checklist is provided as supplementary material.

### Animal Husbandry, Lifespan Analysis

Wild-type *Nothobranchius furzeri* (strain GRZ) were maintained at 27°C on a 12-hour light/dark cycle. Water conditions were controlled (pH 7.0–7.4; conductivity, 2800 µS). For aging studies, fish were housed in a recirculating system; for SGLT2i experiments, fish were maintained in static tanks with daily manual water changes. Adult fish were fed Artemia and Killifeast® or a custom-made Repashy®-based diet from 4 weeks of age. Dapagliflozin was used for SGLT2i analysis. Mating occurred weekly. Lifespan studies were performed in both sexes from 4 weeks onward. Median cohort survival defined the “old” timepoint.

### Histology, Immunostaining, Functional Assays and Quantification

Fish were euthanized with 0.1% MS-222, and kidneys were fixed in 4% PFA, paraffin-embedded, and sectioned at 10 µm. Sections were stained with H&E or used for immunofluorescence (SGLT2 on sections; CD31 on whole-mounts). After blocking with 10% normal goat serum, tissues were incubated with primary antibodies at 4°C, followed by Alexa Fluor–conjugated secondary antibodies. Imaging was performed on Zeiss Axio Observer Z1 or LSM 980 with Airyscan. CD31+ area was quantified using Fiji and normalized to total kidney area. For vascular leakage assays, young and old fish were perfused intracardially with 0.1% fluorescent albumin; kidneys were fixed 48 h later and imaged via spinning-disk confocal microscopy (CSU-W1, Yokogawa), and albumin+ areas were quantified as a percentage of kidney area. For ex vivo tubular transport, kidneys were incubated in marine teleost buffer with fluorescent methotrexate or 2-NBDG, ±MRP2 or SGLT2 inhibitors, and imaged by spinning-disk confocal microscopy; fluorescence intensity ratios (cell/lumen) were analyzed in Fiji. For 3D vascular analysis, fish were perfused with 3% gelatin containing 0.1% fluorescent albumin, kidneys were cleared with formamide, and imaged by LSM 980. Vascular volume, length, density, and branch points were quantified using Fiji, Amira, and WinFiber3D.

### Single-Nucleus RNA Sequencing

Our full single-nuclei isolation protocol is available in supplementary methods. Libraries were prepared using the 10x Genomics Chromium Single Cell 3′ v3.1 platform and sequenced on an Illumina NovaSeq 6000. The single-nucleus RNA-seq datasets generated during this study as well as processed data and code used for analysis are available in the GEO repository under accession number GSE297623. Additional information is available from the corresponding author upon request.

### Statistical Analysis

All data were analyzed in GraphPad Prism. Outliers were excluded using a Q=1% threshold. Normality was assessed via Shapiro-Wilk test. Parametric (unpaired *t* test, one-way ANOVA with Tukey post hoc) or nonparametric (Mann-Whitney, Kruskal-Wallis) tests were used as appropriate. Each data point displayed represents a biological individual; technical replicates based on experimental conditions varied between three to twelve. Data are shown as mean ± SD; significance was defined as *p* ≤ 0.05 (\**), p* ≤ *0.01 (**),* and *p* ≤ *0.001 (\*\*\**).

## Results

### Killifish exhibits advanced aging with stable organ scaling

The African turquoise killifish (*Nothobranchius furzeri*, ATK, henceforth referred to as killifish) is a naturally short-lived vertebrate model, exhibiting rapid growth, early sexual maturation, and a compressed aging timeline (Fig. 1A). The killifish kidney consists of two anatomically distinct regions (Fig. 1B): the head kidney contains glomeruli, tubules, and hematopoietic cells, and the trunk (or tail) kidney comprises primarily collecting ducts. Both kidney regions are covered in pigmented melanocytes, giving them a speckled appearance. In our facility, the median lifespan of the GRZ strain, which is the shortest lived of the killifish strains^13^, is approximately 18 weeks (Fig. 1C). Sexual maturity is reached by 4-6 weeks of age. Sex-specific differences in lifespan are mitigated when animals are housed separately, while group or family housing leads to shorter lifespan in males than females (supplementary Fig. S1A). Killifish exhibit indeterminate growth, with both length and weight increasing—albeit at a reduced rate with age—while body mass index (BMI) remains relatively stable (Fig. 1D). Despite continuous somatic and renal growth, the kidney-to-body weight ratio remains stable, varying only about 0.5-1% across age and sex, suggesting proportional organ scaling during aging (Fig. 1D).

**Figure 1.**
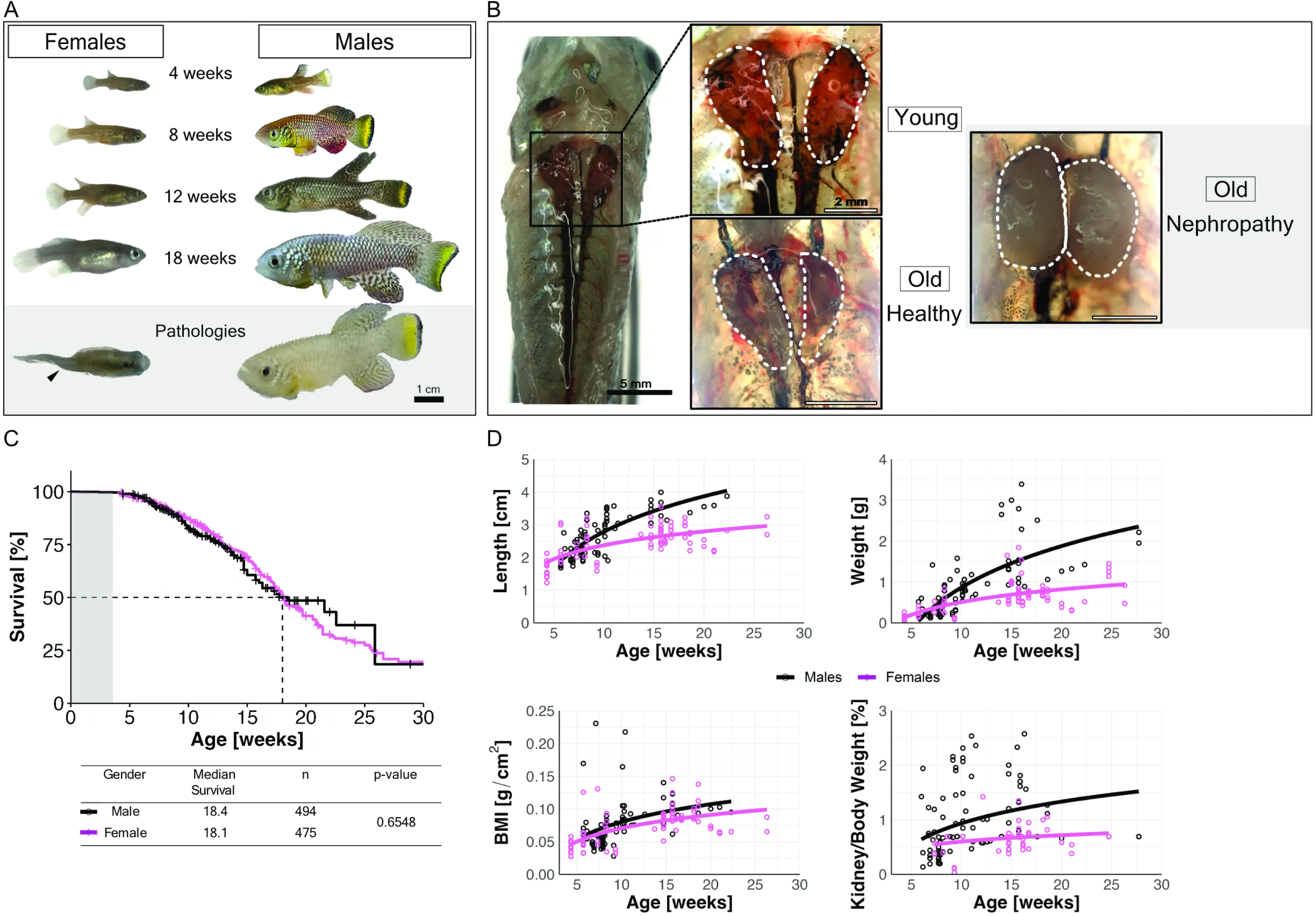
Aging-associated changes and nephropathy in African turquoise killifish. (A) Representative images of female and male killifish at 4, 8, 12, and 18 weeks of age. Lower panel shows phenotypic pathologies in old fish: spinal kinking (left, arrowhead indicates body deformation), dropsy (right). Scale bar: 1 cm. (B) Representative morphology of head kidneys at young and old age. Dotted lines outline head kidney structure. Old fish show examples of healthy kidneys and nephropathy. Scale bars: 5 mm (left); 2 mm (right). (C) Survival curves of male (black) and female (magenta) killifish over time. (D) Growth parameters over time, including body length, body weight, body mass index (BMI), and kidney-to-body weight ratio, stratified by sex. Data points represent individual animals; lines represent fitted regression curves.

### Aging killifish kidney displays hallmarks of nephrosclerosis

Aging in killifish kidneys is characterized by progressive structural deterioration, consistent with hallmarks of nephrosclerosis. Kidney histology reveals a marked contrast between young and aged kidneys (Fig. 2A). In young kidneys, glomeruli, tubules, and vasculature maintain an organized and healthy morphology. In contrast, aged kidneys exhibit an increased proportion of glomerulosclerosis, tubular atrophy, and vascular remodeling, including thickening of larger vessels and the characteristic “onion-layer” atrophy of smaller vessels.

**Figure 2.**
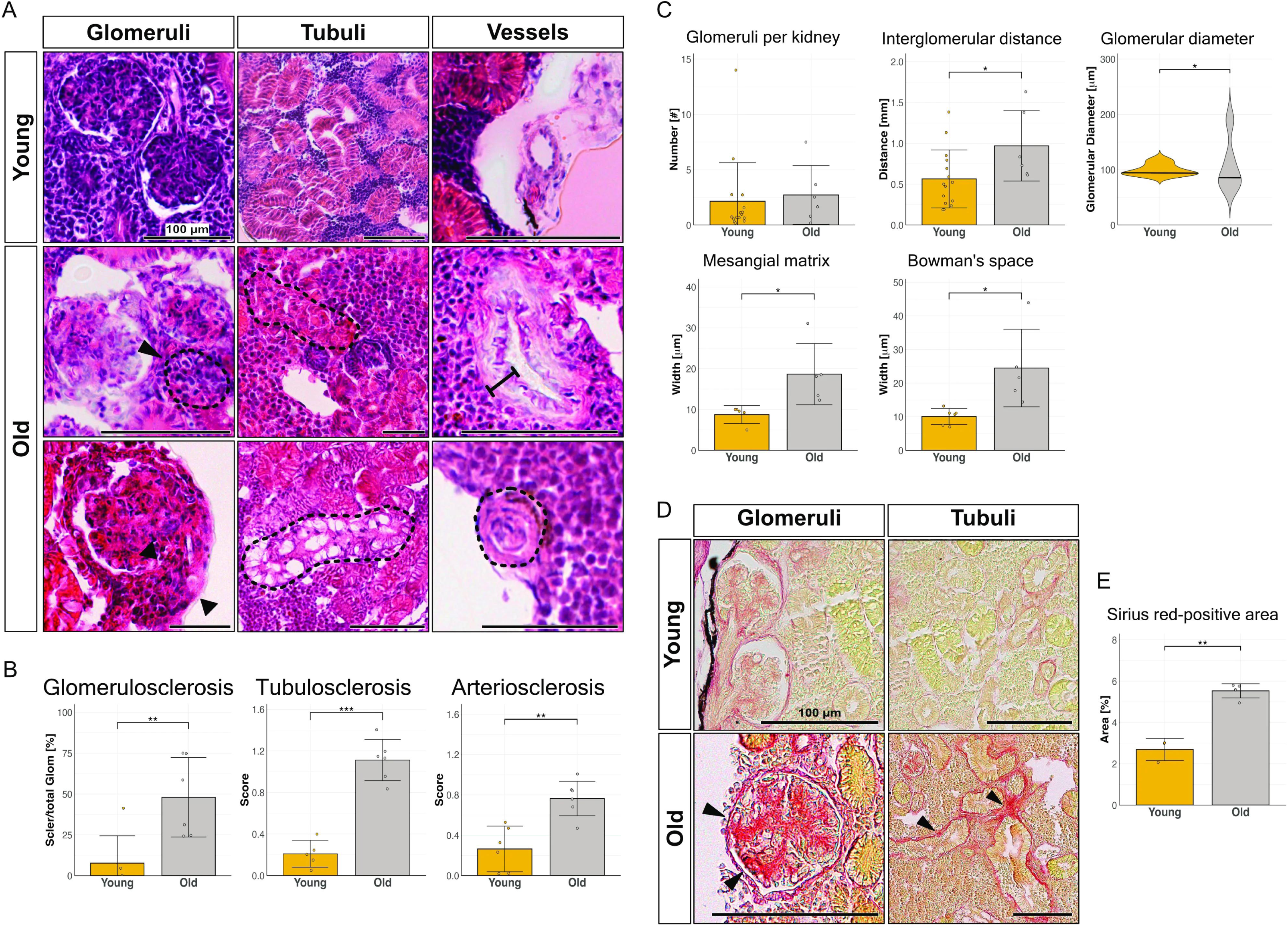
Aging killifish kidneys are characterized by nephrosclerosis and fibrosis. (A) Representative hematoxylin and eosin (H&E) staining of glomeruli, tubuli, and vessels from young (top) and old (bottom) killifish kidneys. Dotted outlines indicate pathological glomeruli and tubuli (e.g., sclerosed glomeruli, tubular atrophy, vacuolized tubules). Black arrowheads point to mesangial matrix thickening. Black lines show vessel wall thickening and “onion-like” atrophic vessels. Scale bar: 100 µm. (B) Quantification of glomerulosclerosis: 7.7 ± 17% in young, 48 ± 24% in old animals (*p = 0.013*), tubulosclerosis (*p = 0.0001*), and arteriosclerosis scores (*p = 0.0015*) (n = 6 young, n = 6 old). (C) Morphometric analysis of glomeruli showing stable glomerular counts but increased interglomerular distance (*p = 0.0266*) suggesting a decrease in glomerular density, greater glomerular size variability with age (*p = 0.0128* using F-test), and mesangial matrix expansion (*p = 0.0219*) and Bowman’s space expansion (*p = 0.0146*) (n = 6 to 16 young, n = 6 old). (D) Representative Sirius Red staining reveals collagen deposition in glomerular and tubular regions (arrowheads). Scale bar: 100 µm. (E) Quantification of Sirius Red–positive area as percentage of total tissue area (n = 3 young, n = 4 old).

Glomerulosclerosis is a key pathological feature of aging in killifish. The proportion of sclerosed glomeruli is about six times greater in old animals than in young animals (Fig. 2B). Mean glomerular diameter remains stable during aging, but the variance between glomerular sizes increases significantly in old kidneys (Fig. 2C). Aged kidneys also show mesangial matrix expansion with significant increase in Bowman’s space (Fig. 2C). Despite no change in the total glomerular count with age, the distance between individual glomeruli increases significantly in aged kidneys (Fig. 2C), suggesting reduced glomerular density with age.

Aging killifish also exhibit tubular degeneration, as tubular atrophy and vacuolization become more prominent (Fig. 2A), and show higher levels of tubulosclerosis (Fig. 2B). Arterioles exhibit pronounced age-related structural changes, including hyalinosis and thickening of the vascular walls, leading to a significant increase in arteriosclerosis (Fig. 2B). Aged kidneys also show increased fibrotic deposition (Fig. 2D, E).

Together, these findings establish the killifish kidney as a model for age-associated nephrosclerosis.

### Killifish develop age-dependent albuminuria and impaired primary active secretion, but preserve glucose uptake

We discovered that after intravascular injection of fluorescently labeled albumin (size: 67 kDa), albumin leaked from the vascular system into tubules, suggesting dysfunctional glomerular barrier integrity. To assess this further, we injected albumin intravascularly into young and old killifish and examined renal accumulation 48 hours post-injection. In young kidneys, minimal albumin retention within the vasculature was observed, with minimal signal observed in renal tubules, indicating an intact filtration barrier. In contrast, aged kidneys exhibited pronounced accumulation of albumin within tubular lumens, consistent with increased glomerular permeability and protein leakage (Fig. 3A, B). These findings indicate an age-dependent decline in glomerular barrier function, manifesting as albuminuria.

**Figure 3.**
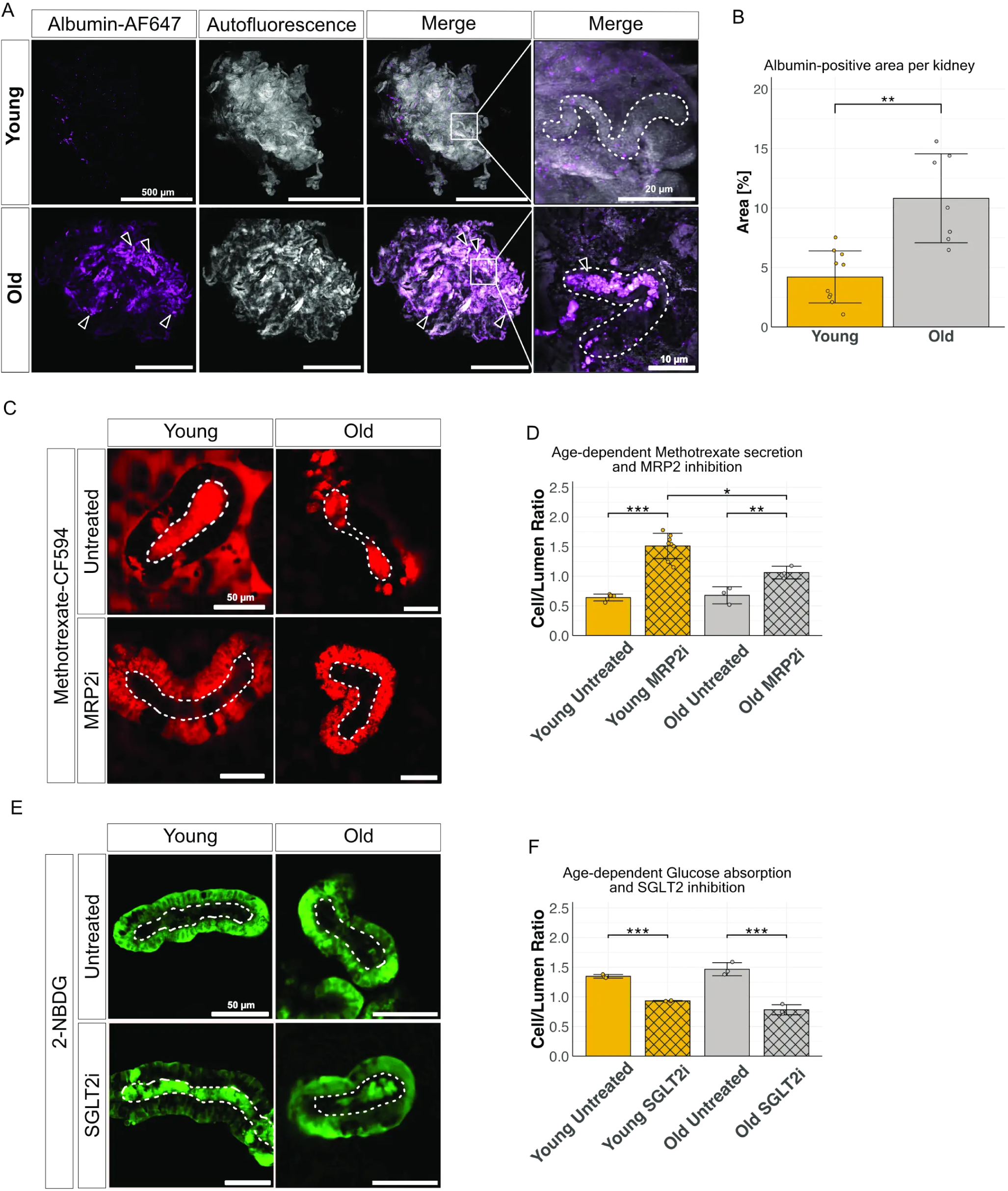
Age-dependent functional changes in the killifish kidney. (A) Representative images of albumin leakage after intracardial albumin injection in young (top) and old (bottom) killifish kidneys. White dotted lines outline tubules, and arrowheads point to tubules filled with albumin in aged kidneys. Scale bars: 500 µm; inset: 10–20 µm. (B) Quantification of albumin-positive area per kidney, showing significantly higher albumin leakage in aged kidneys (n = 10 young, n = 7 old). (C) Representative images of methotrexate (Mtx) secretion in untreated kidneys (top) and kidneys treated with MRP2 inhibition (bottom). Untreated tubules secrete Mtx into their lumen, whereas MRP2i-treated tubules accumulate Mtx intracellularly. (D) Quantification of Mtx secretion (cell/lumen ratio) shows an age-dependent decline in secretion under MRP2 inhibition (*p < 0.01*) (n = 4 young untreated, n = 7 young treated, n = 3 old untreated, n = 3 old treated). (E) Representative images of glucose absorption visualized by 2-NBDG in untreated tubules (top) and treated with SGLT2 inhibition (bottom). 2-NBDG is reabsorbed into the cell: with SGLT2i, 2-NBDG accumulates in the lumen. (F) Quantification of glucose absorption (cell/lumen ratio), showing no significant difference between young and old tubules, nor difference in response to SGLT2 inhibition (n = 3 for all groups).

A physiological change in aging human kidneys is decreased functional drug secretion. We used *ex vivo* analysis of isolated renal tubules to assess age-dependent active drug secretion. Methotrexate (Mtx), a known substrate of the ATP-binding cassette transporter MRP2, was used to evaluate primary active transport. Under normal conditions, Mtx is secreted from tubular cells into the lumen, resulting in a cell/lumen ratio <1 (Fig. 3C, top panel). Pharmacological inhibition of MRP2 reversed this gradient, causing intracellular accumulation of Mtx (cell/lumen ratio >1) (Fig. 3C, bottom panel). While baseline secretion was similar in young and old tubules, the effect of MRP2 inhibition was significantly attenuated in aged samples (Fig. 3D), indicating an age-related decline in ATP-dependent drug transporter activity.

In contrast, proximal tubular glucose uptake remained unaffected by aging. In *ex vivo* assays, fluorescent glucose analog 2-NBDG accumulated in tubular cells from both young and old fish, yielding a cell/lumen ratio >1 (Fig. 3E, top panel). SGLT2i reduced intracellular signal and increased luminal accumulation in both groups (Fig. 3E, bottom panel), consistent with effective transporter blockage. There were no significant age-dependent differences in SGLT2 activity (Fig. 3F), suggesting preserved secondary active glucose reabsorption during aging.

Together, these findings indicate that while aging compromises glomerular barrier integrity and selectively impairs ATP-dependent drug secretion, key solute reabsorption processes such as glucose uptake remain functionally intact in aged killifish kidneys.

### Microvascular rarefaction in the aging killifish kidney

Microvascular rarefaction—the progressive loss of glomerular and peritubular capillaries—is a defining feature of renal aging. We assessed functional vascular density in the killifish kidney via 3D imaging of the vessel architecture. Young kidneys showed an extensive, highly branched vascular network, while aged kidneys displayed visibly reduced vascular complexity (Fig. 4A). Vessel segmentation and quantification confirmed a significant decline in total vessel volume and vessel-to-kidney volume ratio with age. In addition, compared to young kidneys, aged kidneys showed a significant decrease in the ratio of total vascular to kidney volume (Fig. 4B), and in vascular complexity, as reflected in a significant decrease in the total number of vascular branch points Fig. 4B). In addition, aged kidneys showed shorter average vessel length and decreased overall network length (Fig. 4B). Analysis of vessel density based on vessel diameter revealed a specific decline in capillary density—defined by vessel diameter ≤ 7µm—in aged kidneys (Fig. 4C), highlighting how aging affects microvasculature.

**Figure 4.**
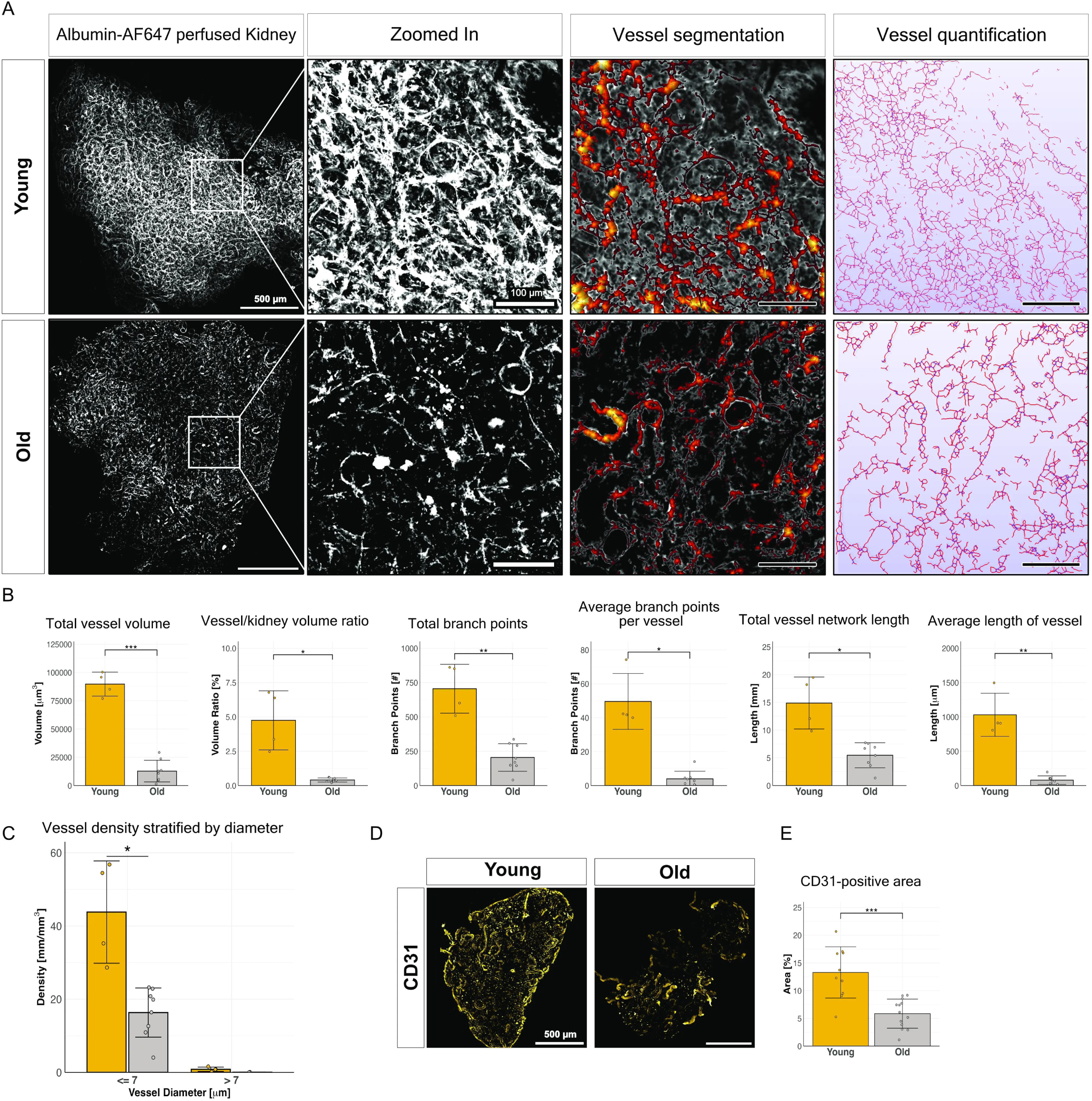
Age-associated changes in vascular morphology and density in the killifish kidney. (A) Representative 3D images of young (top) and old (bottom) killifish kidneys perfused with fluorescently labeled albumin hydrogel. Alterations in vascular structure are more pronounced in old kidneys. The images show a representative maximum intensity projection as well as sections with vessel segmentation performed in Amira and quantification done in WinFiber3D. Scale bars: 500 µm (left); 100 µm (right). (B) Quantification of 3D vessel morphology and volume in young (n = 4) and old (n = 8) kidneys. (C) Bar plot showing vessel density (mm/mm³) stratified by vessel diameter (≤7 μm vs >7 μm) in young (yellow) and old (gray) killifish kidneys. Capillary density is significantly reduced in old kidneys (*p* = 0.0234). (D) Representative whole-mount immunofluorescence images of CD31 staining, in young and old kidneys. Scale bar: 500 µm. (E) Quantification of CD31-positive area relative to total kidney area (n = 12 young, n = 13 old).

We also showed that aged kidneys exhibited a striking reduction of the vascular endothelium, as indicated by immunofluorescence staining with CD31, a pan-endothelial marker (Fig. 4D). Quantification revealed that the CD31-positive area declined by over 50% with age (Fig. 4E). This finding is consistent across sexes (supplementary Fig. S2, top panels), though the relative reduction was more pronounced in males.

These findings establish vascular rarefaction as a conserved and quantifiable hallmark of kidney aging in the killifish.

### Loss of energy homeostasis in killifish kidneys

We next sought to determine which cell populations show a decrease in the kidney with age. We first performed single-nucleus RNA sequencing (snRNA-seq) with young and old male and female fish, using a 10x Genomics Chromium platform. Unsupervised clustering identified 28 distinct cell populations, which were classified based on their transcriptomic profile and then shown via UMAP projection (Fig. 5A). Major renal cell types were represented included podocytes, three proximal tubule subclusters, and distal tubule cells. Cells from the collecting duct could not be confidently annotated and were excluded from further analyses. Non-epithelial populations included endothelial and perivascular cells, with the latter further subclustering into pericytes and myofibroblasts. (See supplementary Fig. S3 for systemic annotations of major renal cell types.) Notably, the most abundant cell populations were derived from the hematopoietic system, comprising B and T lymphocytes, macrophages, and various progenitor populations. In addition, several non-renal cell types were also detected. Cell clusters lacking robust marker gene expression were omitted from downstream analyses.

**Figure 5.**
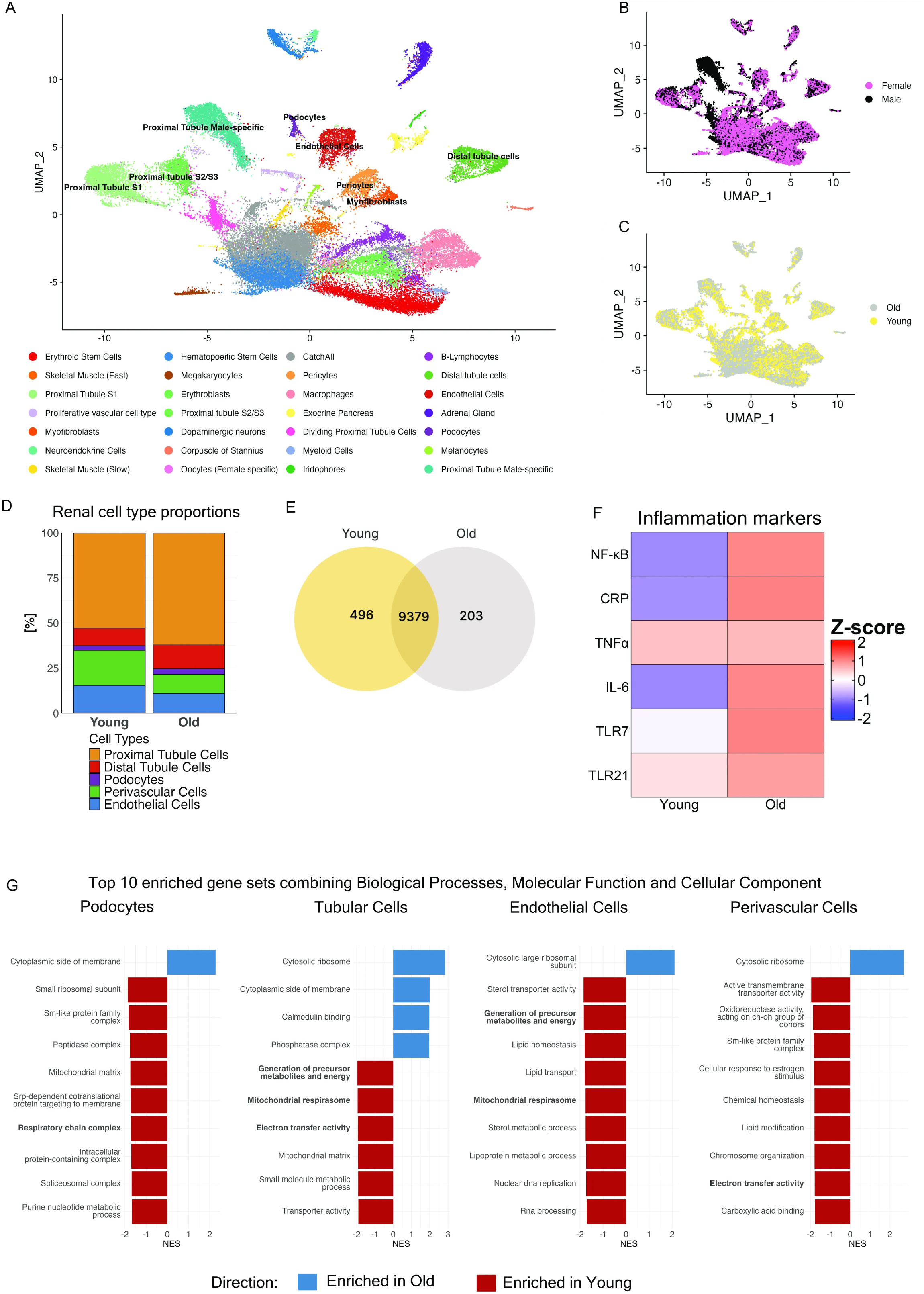
Aging alters renal cellular composition, transcriptional profiles, and energy metabolism in the killifish kidney. (A) UMAP projection of single-nucleus RNA-sequencing data from young and old male and female killifish kidneys reveals 28 distinct renal cell clusters, annotated by cell type. (B) UMAP plot colored by sex shows substantial overlap between male and female transcriptomes, with the exception of a male-specific tubular cell cluster. (C) UMAP colored by age highlights age-associated expansion of hematopoietic populations (age-dependent cell increase by 84%) and reduced abundance of vascular cells. (D) Stacked bar plots showing relative proportions of major renal cell types by age. Endothelial and perivascular cells decline with age (endothelial cells decreased from 15.4% in young to 10.9% in old (-30%) and perivascular cells decreased from 19.4% to 10.7% (-45%) respectively), while proximal and distal tubular cell fractions increase (+17% and +35%), as do podocytes (+16%). (E) Venn diagram displaying differentially expressed genes between young and old kidneys. (F) Heatmap of selected inflammatory markers reveals increased expression of pro-inflammatory genes, including *NF-*κ*B, CRP, TNF-*α*, IL-6, TLR7,* and *TLR21*, in aged kidneys, supporting “inflammaging.” (G) Gene set enrichment analysis of aging-associated transcriptional changes in tubular cells, endothelial cells, perivascular cells, and podocytes. Top 10 enriched gene sets across combined GO categories (Biological Process, Molecular Function, Cellular Component) are shown.

All major cell types were detected across age groups and sexes, with the exception of a subcluster of proximal tubule cells (proximal tubule cluster 3) that was predominantly male-specific (Fig. 5B). Despite this sex-specific distinction, proximal-tubule clusters remained spatially close to each other on the UMAP, whereas distal-tubule populations were clearly segregated.

While the overall clustering structure remained conserved across the two age groups (Fig. 5C), quantitative analysis demonstrated a prominent expansion of hematopoietic populations with age (increase by 84%, data not shown). Within kidney-resident cell types, opposing trends were observed: tubular epithelial cells and podocytes increased in proportion, whereas the proportion of vascular-associated cell types, i.e., endothelial cells and perivascular cells, declined (Fig. 5D). These results suggest that aging in the killifish kidney is accompanied by a shift toward epithelial cell predominance at the expense of vascular and stromal cell populations, consistent with microvascular rarefaction observed in histological and 3D-imaging analyses.

Differential gene expression analysis revealed substantial transcriptional remodeling with age. Of the 10,078 genes detected across conditions, 496 were unique to young kidneys and 203 to old kidneys (Fig. 5E). In addition, a heatmap of canonical inflammatory markers showed upregulation of pro-inflammatory genes in aged kidneys, consistent with a transcriptional signature of chronic low-grade inflammation (“inflammaging”; Fig. 5F).

Gene set enrichment analysis (GSEA) across four major renal cell types revealed consistent downregulation of mitochondrial and energy-related pathways with age (Fig. 5G), supporting an age-associated metabolic shift away from oxidative phosphorylation toward less efficient energy production pathways. Endothelial and perivascular cells showed widespread suppression of lipid transport, sterol metabolism, and oxidoreductase activity, indicating a decline in both metabolic and biosynthetic functions. In endothelial cells a metabolic shift towards glycolysis can be observed with negative enrichment of mitochondrial energy pathway as well as pathways related to lipolysis and amino acid metabolism. In podocytes, negative enrichment was observed for mitochondrial matrix and respiratory chain-related gene sets. Perivascular cells displayed reduced activity in chromatin and chemical homeostasis pathways (see supplementary Fig. S4 for an in-depth overview of the GSEA results).

Together, these findings reveal that aging in the killifish kidney is characterized by loss of mitochondrial energy production and a metabolic shift, particularly affecting vascular compartments, and strongly correlates with structural and functional renal decline.

### Age-associated loss of endothelial cell-cell contacts and intercellular communication

We next investigated age-related changes in intercellular communication in the killifish kidney using CellChat to infer ligand-receptor interactions between renal and vascular cell types. Overall, the total number of ligand-receptor interactions declined with age across all kidney cell types (Fig. 6A), reflecting reductions in cell-cell adhesion, secreted signaling molecules, and extracellular matrix (ECM) components (Fig. 6B).

**Figure 6.**
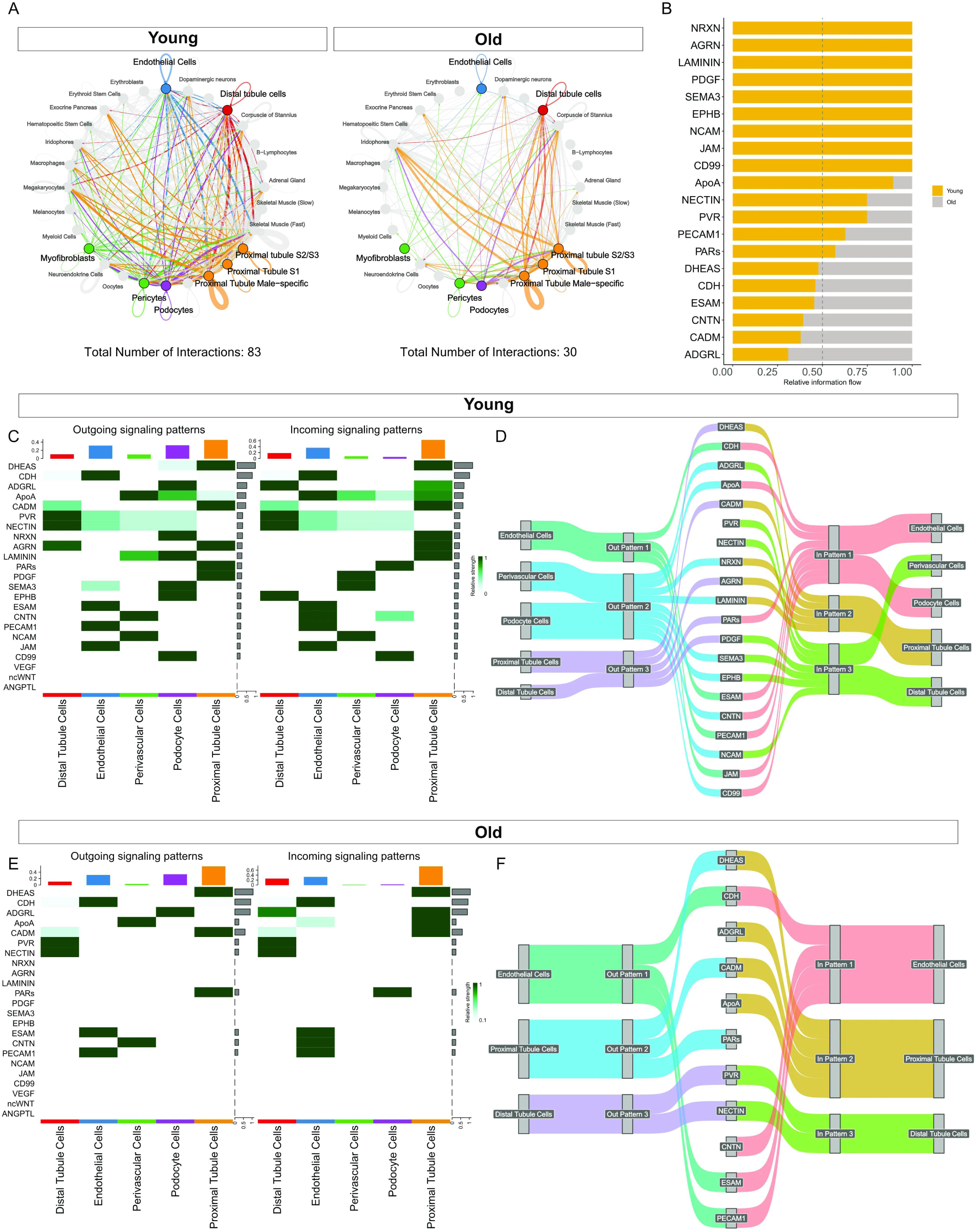
Aging disrupts cell–cell communication networks in the killifish kidney. CellChat analysis was performed on single-nucleus RNA-seq data from kidneys of young and old killifish. (A) Circle plots visualize global intercellular communication networks in young (left) and old (right) kidneys. Each node represents a cell type, and each connecting line a significant interaction, with width proportional to communication strength. Cell types of interest are colored, non-renal cell types displayed in gray. The total number of interactions decreases from 83 in young to 30 in old kidneys. (B) Bar graph shows relative information flow of individual signaling pathways in young (yellow) and old (gray) renal cells, highlighting selective retention or loss of pathway activity with age. (C, E) Heatmaps show outgoing (left) and incoming (right) signaling patterns in young (C) and old (E) kidneys. Pathways are sorted by signaling in young kidneys. Aging reduces both the diversity and magnitude of signaling activities across cell types. (D, F) Sankey plots illustrate dominant outgoing signaling pathways in young (D) and old (F) renal cells, demonstrating diminished pathway complexity and redistribution of signaling roles with age.

A heatmap of signaling patterns in young and old kidneys shows marked loss of intercellular communication with age (Fig. 6C+E). Compared to young kidneys, old kidneys show substantially reduced endothelial cell autocrine signaling, as JAM (junctional adhesion molecules), nectins, and PVR (poliovirus receptor) are lost in old kidneys. Furthermore, old kidneys exhibit reduced signaling of perivascular cells to endothelial cells via CNTN and ApoA, and diminished perivascular ECM signaling, as evidenced by reduced laminin, nectin and PVR signaling. Perivascular cells and podocytes show the strongest age-associated loss of signaling pathways, suggesting an impaired ability of podocytes and perivascular cells to maintain ECM and basement membrane. Together, these alterations indicate a profound disruption of vascular and extracellular homeostasis with aging.

### SGLT2 inhibition preserves renal vasculature and reduces proteinuria

SGLT2 is a key proximal tubular transporter responsible for glucose reabsorption and has emerged as a clinically relevant target in the treatment of diabetic and chronic kidney disease. To evaluate its role in the killifish kidney, we first confirmed the presence of SGLT2 via immunostaining, which revealed apical localization of SGLT2 in proximal tubular cells in aged kidneys (Fig. 7A).

**Figure 7.**
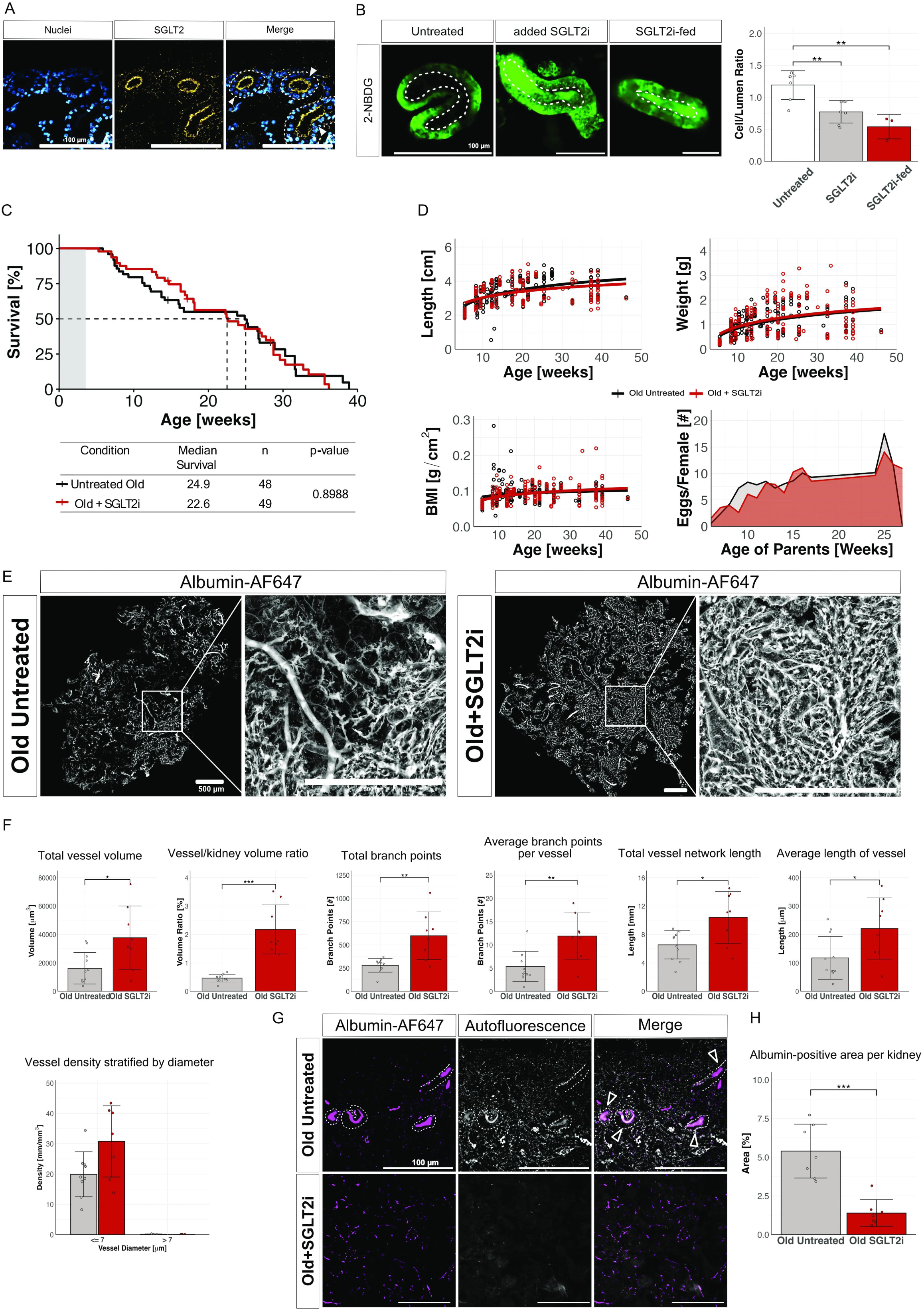
SGLT2 inhibition preserves renal vasculature and reduces proteinuria in aging killifish without extending lifespan. (A) Representative images of kidney section from old untreated killifish stained for SGLT2 (yellow) using a custom-made antibody and nuclei (blue). SGLT2 expression is observed on the luminal side of tubules. Scale bar: 200 µm. (B) Representative image of glucose absorption, visualized by the fluorescent glucose analog 2-NBDG (green), showing decreased glucose uptake in the tubules of old fish kidneys fed with SGLT2i. Quantification of the cell/lumen ratio shows a significant decrease in glucose uptake in old SGLT2i-treated kidneys compared to untreated controls, and no difference between tubules treated directly with SGLT2i *ex vivo* (gray bar) and fish fed SGLT2i (red bar); (n = 7 untreated, n = 7 SGLT2i added directly to tubules, n = 3 SGLT2i-fed). (C) Survival curves comparing untreated and SGLT2i-treated killifish, with median survival at 24.9 weeks for untreated fish and 22.6 weeks for SGLT2i-treated fish. Statistical analysis indicates no significant difference in survival between groups (*p* = 0.8988). (D) Growth parameters of untreated and SGLT2i-treated killifish, including body length, weight, BMI, and fecundity (number of eggs per female). SGLT2 inhibition has a minimal effect on growth or reproductive parameters. (E) Representative images of vessel visualization in old untreated and old SGLT2i-treated kidneys. Kidneys were perfused with albumin. Vessel networks are segmented and quantified, showing differences in vascular structure. Scale bar: 500 µm. (F) Quantification of 3D vessel morphology and volume in old untreated and old SGLT2i-treated kidneys. Total vessel volume, vessel-to-kidney volume ratio, and total branch points are significantly higher in the SGLT2i-treated group. Other vessel parameters, such as average branch points per vessel, vessel network length, and average vessel length, also show significant improvements with treatment. Bar plot shows trend in restoring vessel density (mm/mm³) of vessels with diameter ≤7 μm in untreated (gray) and SGLT2i treated (red) killifish kidneys (*p* = 0.0548), no difference in vessels with diameter >7µm (n = 11 untreated, n = 8 SGLT2i-treated). (G) Representative images of albumin leakage in old untreated and old SGLT2i-treated kidneys. Arrowheads indicate albumin leaked into tubules. SGLT2i treatment reduces albumin leakage in the tubules of aged kidneys. Scale bar: 100 µm. (H) Quantification of albumin-positive area per kidney (n = 7 per group).

We used the SGLT2 inhibitor dapagliflozin in our treatment experiments. To administer dapagliflozin, we developed a custom diet incorporating the drug, thereby enabling long-term pharmacological inhibition while avoiding the substantial problems with water-based delivery, e.g., hydrolysis of dapagliflozin, high drug requirements, uncertain absorption through the gills, and unknown accumulation dynamics. Functional inhibition was verified using an *ex vivo* tubule assay: fish fed the SGLT2i-enriched diet exhibited significantly reduced uptake of the fluorescent glucose analog 2-NBDG compared to fish fed a regular diet (Fig. 7B). The cell-to-lumen ratio was comparable between fish that were fed the drug and tubules treated directly with SGLT2i *ex vivo* (Fig. 7B), confirming effective systemic delivery and target engagement.

We found that although tubules treated directly with dapagliflozin *ex vivo* robustly inhibited glucose uptake (Fig. 7B), dapagliflozin treatment did not extend lifespan in the GRZ strain (Fig. 7C). Sex-specific analysis showed a trend of extended longevity in female fish, but no difference in males (supplementary Fig. S2). SGLT2i treatment resulted in no significant changes in body length, weight, BMI, or fecundity (Fig. 7D).

Strikingly, however, SGLT2 inhibition preserved renal vascular structure in aged animals, as indicated by 3D reconstruction of perfused dapagliflozin-treated kidneys and untreated age-matched controls (Fig. 7E). Quantitative analysis confirmed that SGLT2i resulted in significant increases in total vessel volume, vessel-to-kidney volume ratio, and total branch points (Fig. 7F). However, these effects were far more pronounced in female fish than in males (supplementary Fig. S2). Capillary density showed a trend towards recovery post-treatment, albeit not statistically significant (Fig. 7F). Significance was reached, however, in female kidneys, but not in males (supplementary Fig. S1B).

SGLT2 treatment also partially restored glomerular filtration barrier function. *In vivo* albumin leakage assays revealed a reduction in albumin-positive tubules in SGLT2i-treated kidneys (Fig. 7G), and quantification confirmed a significant decrease in albumin-positive area per kidney (Fig. 7H), suggesting reduced proteinuria in SGLT2i-treated animals. This finding occurred in both sexes (supplementary Fig. S2). In addition, treatment of old kidneys with SGLT2i resulted in an increase in CD31-positive (vascular epithelium) area to almost young-like levels (supplementary Fig. S1C).

Together, these findings highlight the potential of SGLT2i to improve kidney health independently of lifespan extension.

### SGLT2 Inhibition restores mitochondrial function and endothelial communication

We then tested whether treatment could reverse age-associated renal transcriptional changes, by performing snRNA-seq on SGLT2i-treated killifish. Results showed that major renal cell identities were conserved between untreated and SGLT2i-treated animals (Fig. 8A; compare with Fig. 5A). Differential expression analysis showed that while most detected genes were shared, 385 genes were unique to the SGLT2i-treated group, compared to 97 genes in the untreated aged group (Fig. 8B), suggesting a treatment-induced transcriptomic shift. Proportions of vascular cell types did not differ substantially between untreated and SGLT2i-treated kidneys (Fig. 8C), whereas proportions of distal tubule cells increased. Notably, expression of pro-inflammatory markers was attenuated in SGLT2i-treated kidneys (Fig. 8D), indicative of reduced inflammaging.

**Figure 8.**
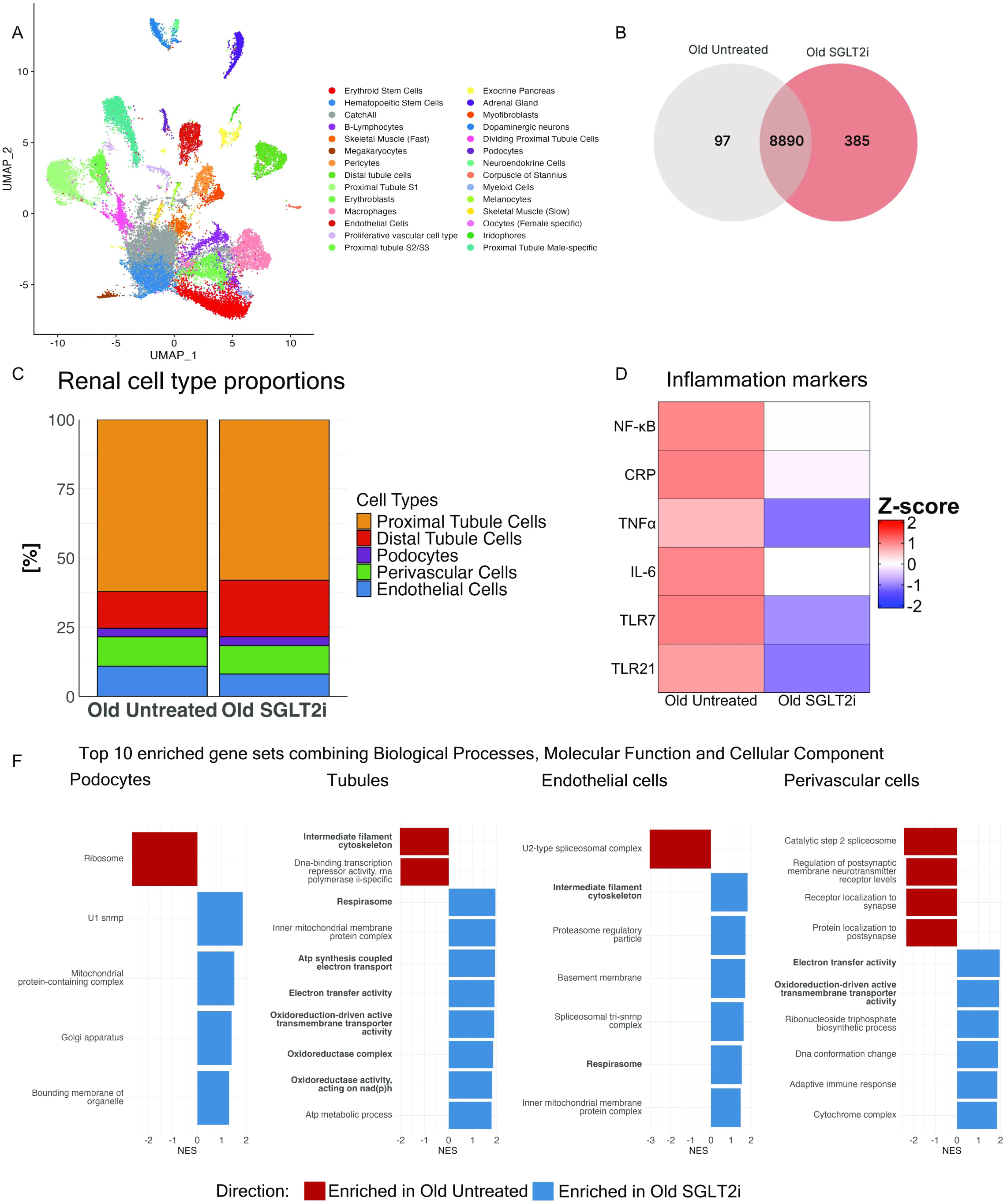
SGLT2 inhibition restores mitochondrial function and reduces inflammatory signaling in aged killifish kidneys. (A) UMAP plot of single-nucleus RNA-seq data from aged SGLT2i-treated killifish kidneys, showing conserved renal cell populations across conditions. (B) Venn diagram depicting shared and condition-specific expressed genes; 385 genes were uniquely detected in SGLT2i-treated aged samples, while 97 were unique to untreated aged samples. (C) Bar plot showing relative proportions of major renal cell types. No major differences were observed in endothelial and perivascular cells counts between the SGLT2i group and the untreated group. (D) Heatmap displaying expression (Z-score) of key inflammatory genes, including *NF-*κ*B, CRP, TNF*α, and *IL-6*, which were downregulated in treated kidneys. (F) Gene set enrichment analysis (GSEA) of major renal cell types showing normalized enrichment scores (NES) for the top 10 gene sets combining biological processes, molecular functions, and cellular components. Positive enrichment (blue) highlights restored expression of mitochondrial and electron transport-associated pathways in SGLT2i-treated animals, particularly in tubular, endothelial, and perivascular cells.

GSEA revealed a striking SGLT2i-mediated restoration of mitochondrial function and cellular energetics across multiple renal compartments (Fig. 8F and supplementary Fig. S5). In endothelial, perivascular, podocyte, and tubular cells, SGLT2 inhibition upregulated pathways associated with electron transport activity, ATP metabolic processes, and mitochondrial membrane integrity. This reversal of age-related suppression in mitochondrial gene sets supports a treatment-induced re-establishment of oxidative phosphorylation and energy homeostasis. In tubular cells specifically, gene sets related to cytoskeletal maintenance and ATP synthesis-coupled electron transport were significantly enriched, suggesting improved structural and metabolic stability.

We also assessed the impact of SGLT2 inhibition on intercellular communication, by performing ligand-receptor network analysis using CellChat (Fig. 9). The numbers of inferred interactions in untreated aged kidneys, which were lower than the numbers in young kidneys, were significantly increased following dapagliflozin treatment (Fig. 9B), particularly among endothelial, perivascular, and podocyte clusters (Fig. 9C). Quantitative interaction strength analysis revealed that SGLT2i preserved key vascular signaling pathways (Fig. 9D). Notably, podocyte-to-endothelium paracrine signaling was enhanced under treatment, characterized by increased VEGFA secretion and activation of endothelial VEGFR2 signaling as well as upregulation of laminin deposition in the ECM.

**Figure 9.**
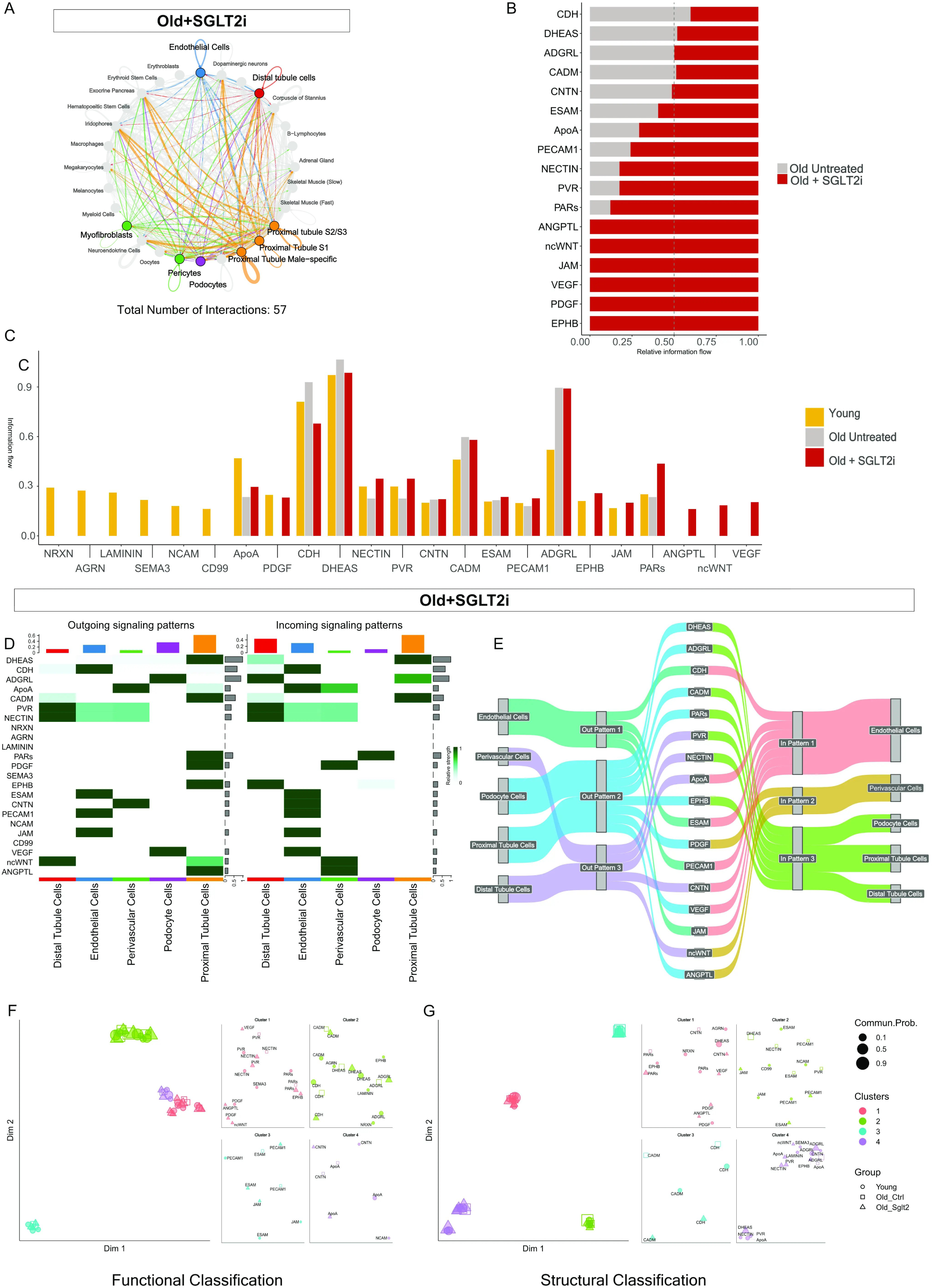
SGLT2 inhibition restores age-related loss of cell–cell communication in the aging killifish kidney. Cell–cell signaling networks inferred using CellChat from single-nucleus RNA-seq data. (A) Circle plot depicts the global intercellular signaling network in kidneys from old fish treated with SGLT2 inhibitor. Each node represents a cell type, and lines represent significant ligand–receptor interactions. The total number of interactions increased to 57 with treatment, compared to 30 in untreated old controls. (B) Bar plot shows relative information flow of key signaling pathways in old control (gray) versus SGLT2i-treated (red) kidneys, indicating selective reactivation of endothelial and perivascular signaling axes. (C) Comparison across all three groups (young, old control, and old + SGLT2i) highlights partial recovery of specific pathways (e.g., PECAM1, JAM, VEGF) toward a youthful signaling profile. Overall, SGLT2i-treated signaling pathways mimic young signaling pathways more than old untreated. (D) Heatmaps illustrate outgoing (left) and incoming (right) signaling patterns in renal cell types from old SGLT2i-treated kidneys. Several signaling axes are re-established, particularly from endothelial and perivascular cells. (E) Sankey plot visualizes major signaling pathways restored in SGLT2i-treated kidneys, with enhanced directional signaling among glomerular and tubular compartments. (F, G) Joint manifold learning was used to classify signaling pathways based on functional (F) and structural (G) similarity across groups. Each dot represents a signaling pathway, clustered by similarity in sender–receiver relationships. SGLT2i-treated kidneys (triangles) show a partial shift toward the youthful communication profile (circles), distinct from untreated aged controls (squares).

Outgoing and incoming signaling patterns confirmed that treated animals maintained robust vascular crosstalk (Fig. 9F). In contrast to untreated aged kidneys, which exhibited collapsed or diminished vascular communication, SGLT2i preserved signaling hubs in endothelial and perivascular compartments. Dimensionality reduction of signaling networks showed that the SGLT2i-treated group clustered more closely with the youthful signature in both functional (e.g., CD99, CNTN, VEGF) and structural (e.g., integrins, cadherins) categories (Fig. 9G–H), highlighting a partial reversion to a regenerative and communicative vascular state.

In addition to preserving age-sensitive vascular signaling pathways such as PDGF, EPHB, and JAM, SGLT2 inhibition uniquely activated a subset of signaling networks not restored in untreated aging kidneys, including ANGPTL, non-canonical WNT (ncWNT), and VEGF. This pattern suggests that SGLT2i does not merely reverse age-related loss of intercellular communication but actively reconfigures the signaling landscape. The emergence of distinct ligand–receptor interactions under treatment points to a drug-specific remodeling effect, highlighting that SGLT2i exerts both rejuvenating and uniquely adaptive influences on renal cell–cell communication during aging.

Together, these results demonstrate that SGLT2 inhibition in the aging killifish kidney restores mitochondrial bioenergetics, reduces inflammatory tone, and preserves endothelial-perivascular communication to a “youthful” state. These effects likely underlie the observed maintenance of vascular integrity despite mild reductions in endothelial cell numbers, offering a mechanistic basis for the vascular-protective effects of SGLT2 inhibition during renal aging.

## Discussion

The African turquoise killifish has emerged as a powerful vertebrate model to study aging across multiple organ systems, including neurodegeneration, immunosenescence, and impaired regeneration. Despite its increasing utility in aging research, renal aging in the killifish has remained largely unexplored. Here, we investigated structural, functional, and transcriptomic changes in the aging kidney and evaluated the effects of SGLT2i as a potential intervention. Using histology, 3D vascular imaging, and single-nucleus RNA sequencing, we reveal hallmarks of nephrosclerosis, progressive microvascular loss and age-associated albuminuria, a metabolic shift from mitochondrial respiration to glycolysis across renal cell types, loss of cell-cell communication and a partial reversal of vascular and metabolic aging features upon SGLT2i treatment.

We observed an increase in interstitial fibrosis in aged kidneys. The main structural hallmark of kidney aging is nephrosclerosis, which is defined by glomerulosclerosis, tubular atrophy, interstitial fibrosis and arteriosclerosis. Our histological analyses revealed that aged killifish kidneys develop classical features of nephrosclerosis closely mirroring those described in aging human kidneys and rodent models ^2,14^. Interestingly, we did not observe (global) glomerular hypertrophy during killifish aging: the mean diameter of glomeruli in aged kidneys stayed similar to that of young animals, but the overall distribution of various sizes increased significantly.

We found an increase in glomerular permeability in aged kidneys through leakage of molecules around 70kDa (like albumin). While albuminuria is not a prominent hallmark of human aging, it is frequently observed in aging rats and certain mouse strains, such as C57BL/6^2^.

We further established age-dependent changes in tubular function. Functional changes observed in mammalian kidneys include changes in tubular reabsorption and secretory capacities, leading to compromised metabolism and clearance of drugs ^14,15^, which can eventually lead to acute interstitial nephritis. We observed diverse age-associated changes in secretion of methotrexate via the MRP2 transporter, an ATP-binding cassette transporter. On the other hand, we were not able to observe differences in SGLT2 function. Isolated proximal tubules of Atlantic killifish (*Fundulus heteroclitus*) have been used as model for pharmacodynamic studies ^16,17^. Atlantic killifish are caught in the wild, and therefore their age cannot be determined. Future studies in the African turquoise killifish will allow for more detailed analysis in tubular transport capacities in an age- and sex-specific manner (Kraus et al, manuscript in preparation).

We found a significant decrease in microvascular density during the aging process. Vascular Rarefaction not only affects vascular density but also endothelial cell integrity and pericyte-endothelial signaling ^18–20^. We observed a decline in CD31-positive staining, reduction of endothelial cell and perivascular cell counts in killifish aging. Consistent with other studies, we observed a ∼45% decline in perivascular cell numbers with age, resembling the 30–50% pericyte loss seen in rodent models of fibrosis post-ischemia^21,22^.

Peritubular capillary loss impairs oxygen delivery, disrupts metabolic homeostasis, and initiates fibrosis and inflammation ^21,23,24^. As such, microvascular rarefaction is increasingly recognized not merely as a consequence but as a driving factor in the progression of renal fibrosis and age-related kidney functional decline ^25–27^. Pronounced microvascular rarefaction was a central finding in aged killifish kidneys, evidenced by loss of capillaries, total vessel volume, reduced vessel length and decline of vessel branching. This is consistent with observations in aged mice, where cortical capillary rarefaction precedes glomerular damage ^28,29^ as well as human kidney transplant recipients, where amount of peritubular capillaries predicts renal function outcome ^30^.

Single-nucleus transcriptomic analysis revealed that aging is associated with widespread downregulation of mitochondrial oxidative phosphorylation and tricarboxylic acid cycle (TCA) cycle pathways across all major renal cell types. The shift from mitochondrial oxidative metabolism toward glycolysis observed in aged killifish kidneys mirrors metabolic inflexibility described in aging mammals ^31^. Concordantly, we observed an age-related loss of ATP-generating pathways and downregulation of fatty acid metabolism and amino acid metabolism gene sets, suggesting impaired metabolic adaptation. Loss of metabolic flexibility also underlies vascular and parenchymal deterioration during aging, in line with recent models reviewed in Augustin & Koh, 2024.

Aged endothelial and perivascular cells exhibited significant loss of cell-cell adhesion molecules, leading to compromised vascular barrier integrity. We further observed an age-associated increase in inflammation markers. This matches a report suggesting that aging and chronic low-grade inflammation progressively destabilize endothelial junctions in other animal models such as mice and zebrafish ^33^. Our findings demonstrate that structural deterioration of cell junctions can occur even in the absence of overt inflammation, emphasizing a direct aging effect and “inflammaging”.

Dapagliflozin treatment restored vascular density, endothelial communication pathways, and reduced albumin leakage in aged killifish kidneys. These effects are consistent with rodent studies showing that SGLT2 inhibitors preserve endothelial function following renal injury^34^. Our study extends this observation to an aging context, suggesting that SGLT2i may be able to partially reprogram endothelial cells. One possible explanation may be the observed upregulation of angiogenic factors such as VEGF and ANGPTL4. These factors were only transcriptionally upregulated in SGLT2i-treated old kidneys. This observation is in line with in aging mice kidneys emphasizing the protective role of VEGF in vascular maintenance and CKD prevention ^21,35–38^.

We also observed that the effects of SGLT2i on vasculature showed sex-dependent differences in vessel length and branching points. A possible explanation could be sex-dependent differences in the pharmacodynamics and -kinetics of SGLT2i also previously described by Miller et al^39^. Despite clear structural and transcriptional improvements with SGLT2 inhibition, we did not observe a significant lifespan extension in treated killifish. This echoes observations from mammalian studies where the benefits of SGLT2 inhibition on vascular and metabolic health do not uniformly translate into lifespan gains, depending on timing and length of drug-intake and sex-specific differences ^39^. The ideal dosing of drugs in a sex-specific manner and the translation of findings in killifish and mice to humans requires further research.

Our study has several limitations. The SGLT2i intervention cohort was relatively small, which may have limited the detection of subtle treatment effects on lifespan. In addition, the fish used in our treatment protocol showed in general an increased lifespan. This could be due to differences in water temperature ^40^ and a difference in diet, highlighting protein source and dietary regime as driver of longevity ^41,42^. Sex-specific data acquisition and analysis was not performed on histological staining as well as tubular assays in this paper. Wherever or not sex-specific analysis was performed, we refer to our supplementary figure S2 for reference.

In summary, we characterized aging kidneys in a novel animal model of aging. We observed an increase in fibrosis, proteinuria and diminished tubular transport mechanisms in aged kidneys. Our study describes important hallmarks of aging ^43,44^ in the kidneys of aging killifish. Mitochondrial dysfunction, including impaired ATP production, inflammation and loss of metabolic flexibility play a central role in aging of all cell types. These changes are associated with altered intercellular communication between endothelial cells, perivascular cells, and podocytes. SGLT2 inhibition not only preserves vascular networks but also reinstates “youthful” transcriptional programs, offering a strategy to mitigate age-associated kidney decline.

## Declaration of Generative AI and AI-assisted technologies in the writing process

During the preparation of this work the authors used ChatGPT 4.0 to improve readability and language of the manuscript. After using this tool, the authors reviewed and edited the content as needed and take full responsibility for the content of the publication.

## Data Sharing Statement

The transcriptomic data supporting the findings of this study are openly available upon publication in the Gene Expression Omnibus (GEO) under accession number GSE297623 (https://www.ncbi.nlm.nih.gov/geo/query/acc.cgi?acc=GSE297623).

*During peer review, the data are accessible to editors and reviewers using a secure token, which is available upon request from the corresponding author*.

All raw and processed imaging data are available via our institutional OMERO server: https://omero-pub.mdibl.org/pubs/paulmann-et-al-2025

## Acknoweldgements

Research reported in this publication was supported by an Institutional Development Award (IDeA) from the National Institute of General Medical Sciences of the National Institutes of Health under grant numbers P30GM154610 and P20GM103423.

Image collection, processing, and analysis for this manuscript were performed with the assistance of **Dr. Frederic Bonnet** and **Hannah Somers** at the MDI Biological Laboratory Light Microscopy Facility (RRID:SCR_019166), supported by the IDeA program under grant number P20GM103423.

We thank the MDI Biological Laboratory Animal Core Facility, particularly Animal Core Director **Edward Seckler**, for assistance with the design and preparation of experimental fish food, and **Gwendolyn Joy** for her exceptional care of the experimental animals.

We gratefully acknowledge the following individuals and groups for their valuable contributions:

The Drummond Lab at MDIBL, especially **Dr. Iain Drummond**, **Dr. Caramai “Nana” Kamei**, and **Will Sampson**, for their support with molecular biology methods in fish models.

The Murawala Lab, especially **Dr. Prayag Murawala**, **Dr. Marco Pende**, **Dr. Sofia-Christina Papadopoulos**, **Vijayishwer Jamwal**, and **Erik Figura**, for their assistance in single-cell and single-nuclei preparation, protocol development, and tissue clearing for imaging.

**Roy McMorran**, Director of Information Technology at MDIBL, for his dedicated support with software access, data management, and IT infrastructure throughout this project.

**Dr. James Godwin**, for his help in designing custom-made antibodies.

**Christine Smith**, supervisor of the DNA and Gene Expression Core Facility, for her support during the establishment of our single nuclei sequencing pipeline.

**Dr. Gert Fricker** and **Lisa Kraus**, for generously sharing their extensive experience and teaching of the killifish tubule assay, a method Dr. Fricker has employed at MDIBL for nearly four decades.

The students and participants of the Origins of Renal Physiology for Medical Students: TREKS 2024 and Origins of Renal Physiology: Fellows 2024 courses, who replicated many experiments described in this study. We further thank the course organizers, Dr. Mark Zeidel, Dr. Bryce McIver, and proximal tubule module leaders Dr. Stewert Lecker, Dr. Moe Orson, and also Dr. Michael Romero, for their insightful discussions, support, and commitment to advancing physiology education in non-canonical model organisms.

This work was supported by funding from the McKenzie Foundation, Morris Discovery Fund, Mount Desert Island Biological Laboratory, and AstraZeneca.

**Acknowledgment of Prior Abstract Publication:**

A preliminary version of this work was presented in abstract form at the American Society of Nephrology Kidney Week 2024. Abstracts are not considered prior publication. ^45^

Paulmann A, Cox M, Beverly-Staggs LL, Johnson CP, Somers H, Haller H. Loss of Kidney Microvasculature during Aging Is Dependent on Several Pathways: A Study in African Turquoise Killifish: TH-PO943. *J Am Soc Nephrol*. 2024;35(10S):10.1681/ASN.2024hh7ttpp2. doi:10.1681/ASN.2024hh7ttpp2

---

**Supplemental Figure S1. Additional graphs showing comparison between all three groups**

(A) Survival curve of family-housed male and female killifish shows a sex-specific age differences compared to single housing (log-rank test, *p* = 0.0349).

(B) Sex-specific vascular density (vessel diameter <7 μm) in old animals treated with SGLT2i (mean ± SD, p < 0.01, *p < 0.001).

(C) Quantification of CD31-positive vascular area in young, untreated old, and SGLT2i-treated old kidneys (p < 0.01, *p < 0.001).

(D) Renal cell type proportions across groups based on snRNA-seq.

(E) Heatmap of inflammatory gene expression (Z-scores) in renal tissue.

(F) Quantification of total interactions per kidney across age groups.

**Supplemental Figure S2. Sex-specific analysis of CD31-positive area and SGLT2i effects**

(A) CD31-positive vascular area in young and old female and male kidneys, with and without SGLT2i treatment (*p* < 0.05, p < 0.01).

(B) Survival curves of female and male killifish treated with SGLT2i versus control (log-rank test, *p* < 0.05).

(C) Quantification of vascular parameters in 3D-reconstructed kidneys: total vessel volume, vessel-to-kidney volume ratio, total branch points, total vessel network length, and average vessel length per branch point, separately for females and males.

(D) Albumin-positive area as a measure of proteinuria in female and male kidneys (mean ± SD, *p* < 0.05, p < 0.01).

**Supplemental Figure S3.**

Dot plot of marker gene expression across renal cell types in killifish kidneys.

**Supplemental Figure S4.**

REVIGO gene ontology enrichment comparing young and old killifish kidneys. Tree maps display enriched GO terms by category (Biological Process, Molecular Function, Cellular Component) in podocytes, tubular, endothelial, and perivascular cells of male and female fish. Colors indicate normalized enrichment scores (NES); red denotes enrichment in old, blue in young.

**Supplemental Figure S5.**

REVIGO gene ontology enrichment comparing untreated and SGLT2i-treated old killifish kidneys.

Tree maps show differentially enriched GO terms across renal cell types and sexes, categorized by Biological Process, Molecular Function, and Cellular Component. Red indicates enrichment in SGLT2i-treated samples, blue in untreated controls.

