## Supplementary Methods for "SGLT2 Inhibition Ameliorates Age-Dependent Renovascular Rarefaction"

### Animal Husbandry

All the work reported here was performed in wild-type African turquoise killifish (*Nothobranchius furzeri*) of the strain GRZ (origin: Gona-Re-Zhou National Park). All procedures were approved by the MDIBL Institutional Animal Care and Use Committee (IACUC, #24-02) and complies with the MDI Biological Laboratory Institutional Assurance # D16-00341. Our husbandry protocol was adapted from Sandhi et al, *manuscript in preparation*.

Fish were maintained at 27°C in an Aquaneering® recirculating system under a 12h light/dark cycle. Water parameters were continuously monitored (pH 7.0–7.4, conductivity 2800 µS). Adult fish were housed in age-matched groups, with adult males in 3 L tanks adjacent to 2-4 females in 6 L tanks and mated once weekly. Based on tank and group size, plastic plants (Aquaneat®) or igloos (Plexx®, Catalog #13220B) were added for environmental enrichment. Embryos were collected, disinfected in 550ppm PVP Iodine (Ovadine®, Syndel, USA), and incubated at 28°C in Ringer’s Solution. At the black-iris stage (12-14 days post fertilization (dpf)), embryos were transferred to coconut plates with Whatman® Filter paper and stored at 27°C for three weeks. Golden eyed embryos were hatched using a salt solution (1g Instant Ocean® in 1L RO water, pH=6, 4°C). Hatchlings were reared at controlled densities and transitioned to system water at 7 days post-hatch (dph). Fish were fed Artemia and Killifeast® (Brine Shrimp Direct, USA) according to age-specific schedules. Under these conditions, the median lifespan was 18 weeks for both sexes. Group housing males and females led to a relative decrease in male lifespan in our facility (female median lifespan: 20 weeks, male median lifespan: 16 weeks, data now shown). "Young" and "old" time points were defined as 6–8 weeks, where sexual maturity was reached, and 18 weeks, respectively.

Experimental fish for the SGLT2 inhibitor trial were housed separately from the circulating system. Males and females were group-housed in a 6 L tank with 1 male and 2-3 females. Large cohorts hatched on the same day were split in a 1:1 ratio to a control and interventional group at the age of 4 weeks.

#### Experimental Food Recipe

Experimental diet consisted of custom-made food. 200ml of reverse osmosis water was brought to a boil in a microwave. 3 teaspoons (tsp) of Repashy® Grub *Pie Insectivore Gel Premix* (Repashy® Speciality Pet Products, USA) and ¾ tsp of Repashy® Redrum *Carotenoid Gel Premix* (Repashy® Speciality Pet Products, USA; for additional food coloring) were added, immediately stirred and let to cool to ~45°C. The liquid premix was poured onto aluminum-covered trays as a thin layer and let to set at room temperature. For drugged food, the SGLT2 inhibitor Dapagliflozin was added for a final drug concentration of 2µM. DMSO was added to Control food; final concentration of DMSO in both foods was 0.1% v/v. After gel solidified into thin sheets, these were dehydrated at 45°C for 12 hours in a food dehydrator (Septree^TM^, USA). Once completely dried, sheets were cut into small flakes using a Slap^TM^Chop (USA). Fish were fed 3x times daily with a spoon, with approximately 60mg food/fish/day, resulting in a SGLT2i dosage of approximately 0.04mg/kg fish/d (equivalent to 3mg Dapagliflozin daily for a 70kg person). Diet was supplemented with Artemia once daily. Water changes were performed daily with fresh system water and the remaining food was cleaned out. Dosages of equivalent to 10mg Dapagliflozin/d/70kg led to a decrease in lifespans compared to control diet, therefore a lower concentration was chosen for downstream experiments.

#### Lifespan Analysis

Lifespan analyses were conducted on both sexes, with mortality recorded daily from week 4 onward. Kaplan-Meier survival curves were analyzed using the log-rank test in GraphPad Prism. Lifespan data includes all data collected from our colony from October 2023 to March 2025. A total of 968 animals were analyzed (494 males, 475 females). Death was censored if animals were used for experiments; censorship rate was 82% in males and 62% in females. Total number of natural death events was 273.

Experimental lifespan data includes data collected from 3 cohorts born between February 2024 and December 2024. A total of 48 animals were in the SGLT2i group (32 males, 17 females), 49 animals were in the control group (30 males, 19 females). Censorship rate in the Control group was 24%, in the SGLT2i group 20%. Total number of natural death events was 73.

#### Growth and weights measurements

Growth assessments included weight and length measurements at specified time points. Fish were anesthetized using MS-222 (Syncaine®, Syndal,USA) 0.2g/L (equivalent to 0.02% w/v, pH 7.2) for 2 minutes, blotted dry, and weighed on an Adventurer™ Pro Analytical Electronic Balance (model: AV64C, Ohaus®, USA).

Images for length measures were either taken using a Zeiss Axio Zoom.V16 (refer to Microscopy section) or using a cellphone, with subsequent transfer of image data in .png format. Length measurements were performed using FiJi (v1.54) ^1^ with calibrated pixel dimensions (1cm = 1541pixels at 10x magnification). Body length was measured manually from the tip of the snout to the end of the tail fin using the line tool.

Kidney weights were determined after euthanizing fish using immersion in MS-222 at 0.5g/l (equivalent to 0.05% w/v, pH 7.2) for 10-15 minutes. Gill movements completely ceased, and as secondary measure the head is cut off using a blade. Head kidney and tail kidney were removed and measured using the above-mentioned scale.

### Histology

Fish were euthanized as described above. Kidneys were fixed in 4% paraformaldehyde (PFA) in phosphate buffered saline (PBS) and kept rocking at 4°C overnight, dehydrated in an ethanol-series and embedded in paraffin. Sections (10 µm) from young and old fish were cut using a microtome (Jung Supercut® Model 2065, Leica, USA) and stained with hematoxylin and eosin (H&E) using standard procedures (H&E Stain Kit, Vector Laboratories®, USA).

### Albuminuria Assay

Killifish were anesthetized using MS-222 (Syncaine®, Syndal,USA) 0.2g/L (equivalent to 0.02% w/v, pH 7.2) for 5 minutes, until gill movements slowed down (but did not cease). Fish were perfused with fluorescently labeled albumin from BSA conjugated with AlexaFluor 647 (Thermo Scientific, Catalog #A34785) at a concentration of 0.1% w/v in phosphate buffered saline (PBS) after intracardial injection following a protocol modified from Mariën et al., 2022. The thoracic cavity was opened with surgical scissors and the beating heart exposed. A microinjection needle was inserted into the heart and 300-500µl of albumin were slowly injected over 3-5 minutes. Successful injections could be visibly monitored by a color change in the heart. Fish were returned into their respective tanks with fresh water and monitored for two days. Two days following the injection the fish were sacrificed, and kidneys were collected. The kidneys were then fixed in 4% PFA for 4 hours at room temperature. They were then washed 3 times for 30 minutes, and stained with DAPI (5µg/ml) for 15 minutes. Kidneys were mounted with Vectashield® Plus Antifade Mounting Medium (Vector laboratories) and a No. 1.5 cover slip (Gold Seal Cover Glass, Thermo Scientific, USA).

### Kidney Tubule Ex vivo Measurements

We used a modified protocol previously published in Atlantic killifish (Fundulus heteroclitus) (^3,4^ to measure ex vivo tubular transport activity. Data displayed was collected in male killifish.

Killifish were sacrificed and cut along the belly with sharp small scissors and guts were removed. Head kidneys were carefully transferred with fine forceps to a dish containing marine teleost saline buffer (Forster Buffer): 140 mM NaCl, 2.5 mM KCl, 1.5 mM CaCl_2_, 1.0 mM MgCl_2_ and 20 mM Tris base at pH 8.0 (Forster and Taggart, 1950). Fine forceps were used to isolate proximal tubules from surrounding connective tissues and hematopoietic cells. The isolated kidney tubules were transferred to tissue culture dishes with cover glass bottoms (FluoroDish®, dish Ø 35 mm) containing 1.0 ml Forster Buffer. All tissue isolations were performed at room temperature (24°C). After isolation, tubules for inhibitor analysis were incubated with respective inhibitors, MK571, MRP2 inhibitor (Selleck, USA, Catalog No. S8126), or Dapagliflozin, SGLT2 inhibitor (Selleck, USA, Catalog No.S1548) for 30 minutes at a final concentration of 2µM and kept in the dark in an aluminium covered dish. Following this, a corresponding fluorescent substrate was added at a concentration of 2µM for 60 min to reach steady-state distribution; either Methotrexat-CF 594 (Biotium, USA, Catalog #00093) or 2-NBDG (2-N-7-Nitrobenz-2-oxa-1,3-diazol-4-ylAmino-2-Deoxyglucose, Catalog #N13195, Thermo Scientific, USA) as glucose analog.

### CD31 Immunofluorescence Labeling

For CD31 staining whole-mount kidney killifish were intracardially perfused using PBS and 4% PFA (200-800µl, while heart was still pumping), after which kidneys were dissected and post- fixed 4% PFA overnight. Samples were blocked in 10% normal goat serum serum (Vector Laboratories), incubated with primary anti-CD31 (1:100, Human CD31/PECAM-1 Antibody, Catalog #: AF806, R&D systems) for three days at 4°C, followed by secondary antibody labeled with Alexa Fluor 647 (Goat Anti-Sheep- AF647, Catalog # A-21448, Thermo Scientific, USA). Kidneys were cleared with a formamide series (20% - 30min, 40% - 30min, 80% - 2h, 95% - 2h twice all at room temperature and gentle rocking) and mounted with Vectashield® Plus Antifade Mounting Medium (Vector laboratories) and a No. 1.5 cover slip (Gold Seal Cover Glass, Thermo Scientific, USA).

### 3D Analysis of Vessel Networks

Additionally, to CD31 staining, vessels were measured in 3D after perfusion with an albumin hydrogel. Protocol was modified and adapted from Lugo-Hernandez et al., 2017.

Male and female killifish were anesthetized using MS-222 (Syncaine®, Syndal,USA) 0.2g/L (equivalent to 0.02% w/v, pH 7.2) for 5 minutes, until gill movements slowed down (but did not cease). Fish were perfused with a hydrogel made from 3% gelatin in Phosphate buffered saline with 0.1% w/v fluorescently labeled albumin from BSA conjugated with AlexaFluor 647 (Thermo Scientific, Catalog #A34785). Immediately after intracardial injection fish were submerged in an ice-bath for 15-20 minutes for the hydrogel to solidify. Kidneys were isolated and fixed in 4% PFA for 24 hours at 4°C with gentle rocking. Tissues were cleared in a formamide series (20% - 1h, 40% - 1h, 80% - 2h, 95% - 2h all at room temperature and gentle rocking, second 95% - overnight at 4ºC), mounted with Aqua-Poly/Mount (Polysciences, USA) on a Superfrost® slide (Thermo Scientific, USA) and a No. 1.5 cover slip (Gold Seal Cover Glass, Thermo Scientific, USA).

### Microcopy methods

All raw and processed microscopy images used in this study, including immunofluorescence and vascular reconstructions, are accessible on our institutional OMERO server at: <https://omero-pub.mdibl.org/pubs/paulmann-et-al-2025>.

#### Whole animals images and length measurements

Whole animal images (used in Figure 1) were acquired using a Zeiss Axio Zoom.V16 (ref: 435080-9031-000, Carl Zeiss Microscopy, Germany) equipped with a Zeiss Plan Z 1.0x/0.25 objective lens (ref: 435282-9100-000, Carl Zeiss Microscopy, Germany), a transillumination base 300 (ref: 435533-9500-000, Carl Zeiss Microscopy, Germany) equipped with a transillumination top 450 mot (ref: 435500-9000-000, Carl Zeiss Microscopy, Germany) in Brightfield mode, and a mechanical stage 150*100 Mot (ref: 435465-9000-000, Carl Zeiss Microscopy, Germany) equipped with an insert plate S, glass 237x157x3 mm ( ref:435465-9053-000 Carl Zeiss Microscopy, Germany). Images were acquired with a Zeiss Axiocam 506 color camera (ref: 426556-0000-000, Carl Zeiss Microscopy, Germany) controlled with Zen 3.1 (Carl Zeiss Microscopy, Germany), at zoom 0.7x, binning 1.1, at resolution of 7740x6249 pixels, in 42 bit and saved in CZI format.

#### Hematoxylin and eosin images

H&E Images were acquired using a widefield Zeiss Axio Observer.Z1 inverted microscope (ref: 431007-9902-000, Carl Zeiss Microscopy, Germany) equipped with an EC Plan-Neofluar 40x/0375 M27 objective lens (ref: 420360-9900-000, Carl Zeiss Microscopy, Germany). Samples were illuminated with a Visible LED (ref: 423053-9030-000, Carl Zeiss Microscopy, Germany), through a Polarizer D (ref: 000000-1121-813, Carl Zeiss Microscopy, Germany), a LD condenser (ref: 424244-0000-000, Carl Zeiss Microscopy, Germany) equipped with DIC prism II/0.55 (ref: 000000-1005-867, Carl Zeiss Microscopy, Germany) and the Analyzer module Pol ACR P&C for transmitted light (ref: 424937-0000-000, Carl Zeiss Microscopy, Germany). Images were acquired with a Zeiss Axiocam 305 color (Ref: 426560-9030-000, Carl Zeiss Microscopy, Germany) controlled with Zen Pro 3.1 (Carl Zeiss Microscopy, Germany) software, at no zoom, binning 1.1, at resolution of 6693 x 9828 pixels, 36 bit depth and saved in CZI file format.

#### Albuminuria Assay

Whole mount kidney images were acquired using a Spinning-disk confocal unit (CSU-W1, Yokogawa, Japan) on a Nikon inverted Ti-Eclipse microscope stand (Nikon Instruments Inc., Japan), equipped with a CFI Plan-Apochromat λ D 4x/0.2 objective lens (ref: MRD70040, Nikon Instruments Inc., Japan)

DAPI, Autofluorescence and Alexa-Fluor 647 were excited with 405 nm (50mW), 488 nm (60mW) and 594nm (40 mW) at 100% intensity from a LUNF-XL laser combiner (77098033, Nikon Instruments Inc., Japan) and collected using Yokogawa DM 405/488/561/640 (ref: MHE46420) with BrightLine® quad-band bandpass filter – 440/521/607/700 nm (ref: FF01-440/521/607/700, Semrock). Z-stack images were collected with a step size of 15.8µm with the Scanning stage (ref: Ti-S-ER, Nikon Instruments Inc., Japan) and the GS35-M stage insert.

Images were acquired in 2048x2048 pixels, Binning 1x1, Pixel size: 1.625µm, in 16-bit with a Scientific CMOS Zyla 4.2 (Andor Technology, United Kingdom) controlled with NIS AR 5.41.02 (build 1711, Nikon Instruments Inc., Japan) software and saved in Nd2 file format.

#### Kidney tubule ex vivo

Tubular images were acquired using a Spinning-disk confocal unit (CSU-W1, Yokogawa, Japan) on a Nikon inverted Ti-Eclipse microscope stand (Nikon Instruments Inc., Japan), equipped with a CFI Plan Apochromat Lambda 20x/0.75 objective lens (ref: MRD00205, Nikon Instruments Inc., Japan).

Mtx-CF594 and 2-NBDG were excited with 594nm (40 mW) and 488nm (60mW) at 100% intensity from a LUNF-XL laser combiner (77098033, Nikon Instruments Inc., Japan) and collected using FF459/526/596-Di01 or dichoric mirror DM 445/514/594 (ref: 99226, Nikon) with 525/50 nm BrightLine® single-band bandpass filter (500 – 550 nm, ref: FF01-525/50, Semrock). Z-stack images were collected with a step size of 1.8µm with the Scanning stage (ref: Ti-S-ER, Nikon Instruments Inc., Japan) and the GS35-M stage insert.

Images were acquired in 1024x1024 pixels, Binning 2x2, Pixel size: 0.6431µm, with a Scientific CMOS Zyla 4.2 (Andor Technology, United Kingdom) controlled with NIS AR 5.41.02 (build 1711, Nikon Instruments Inc., Japan) software and saved in Nd2 file format.

#### Immunofluorescence Imaging

Whole kidney images for CD31 staining were acquired with a Zeiss LSM 980 confocal microscope (Carl Zeiss Microscopy, Germany) on a Zeiss Axio Examiner Z1 upright microscope stand (ref: 409000-9752-000, Carl Zeiss Microscopy, Germany) equipped with a Plan-Apochromat 10x/0.45 M27objective lens (ref: 420640-9900-000, Carl Zeiss Microscopy, Germany). Alexafluor 647 fluorescence was excited with the 639nm line at 1-2% intensity from a 25-mW laser diode and collected using a Airyscan 2 GaAsp PMT detector (ref: 000000-2183-168) with detection wavelengths from 499 to 557 nm and 659 to 720 nm and a BP 570-620 nm + LP 655 filter. Images were sequentially acquired in Super Resolution mode (SR) at zoom 1.7, with a line average of 1, a resolution of 9167 x 7846 pixels, 0.346x0.346x1.740 µm pixel size, a pixel time of 0.73 µs, in 16-bit, and in bidirectional mode. Z-stack images were collected with a step size of 2.9 µm with the Motorized Scanning Stage 130 × 85 PIEZO (Carl Zeiss Microscopy) mounted on the Z-piezo stage insert WSB 500 (ref:0000000-2248-929, Carl Zeiss Microscopy, Germany). The microscope was controlled using Zen Blue Software (Zen Pro 3.1), Airy scan images were processed manually in 3D auto mode with a strength of the deconvolution set to 3.4 and saved in CZI format. All imaging parameters were identical between experiments for compared image.

### 3D vessel imaging

Whole kidney images were acquired with a Zeiss LSM 980 confocal microscope (Carl Zeiss Microscopy, Germany) on a Zeiss Axio Examiner Z1 upright microscope stand (ref: 409000-9752-000, Carl Zeiss Microscopy, Germany) equipped with a Plan-Apochromat 10x/0.45 M27objective lens (ref: 420640-9900-000, Carl Zeiss Microscopy, Germany). Alexafluor 647 fluorescence was excited with the 639nm line at 1-2% intensity from a 25-mW laser diode and collected using a Airyscan 2 GaAsp PMT detector (ref: 000000-2183-168) with detection wavelengths from 499 to 557 nm and 659 to 720 nm and a BP 570-620 nm + LP 655 filter. Images were sequentially acquired in Super Resolution mode (SR) at zoom 1.7, with a line average of 1, a resolution of 9167 x 7846 pixels, 0.346x0.346x1.740 µm pixel size, a pixel time of 0.73 µs, in 16-bit, and in bidirectional mode. Z-stack images were collected with a step size of 1.7 µm with the Motorized Scanning Stage 130 × 85 PIEZO (Carl Zeiss Microscopy) mounted on the Z-piezo stage insert WSB 500 (ref:0000000-2248-929, Carl Zeiss Microscopy, Germany). The microscope was controlled using Zen Blue Software (Zen Pro 3.1), Airy scan images were processed manually in 3D auto mode with a strength of the deconvolution set to 3.4 and saved in CZI format. All imaging parameters were identical between experiments for compared images.

### Image Analysis methods

#### Histology and pathology quantifications

H&E images were analyzed from 6 young and 6 old (male) fish, analyzing 10-15 representative sections per individual fish and averaging the results. Glomerular counts were additionally performed on 10 young fish (n=16 young).

Glomerulosclerosis was calculated by counting the number of visible sclerosed glomeruli and dividing by the number of total glomeruli. The glomerular diameter was measured from vascular pole to the opposite end of the glomerular tuft. Distance between glomeruli was measured from the center of a glomerulus to its nearest neighbor. Mesangial width and space were measured at three points for each glomerulus, directly opposite the vascular pole at 6 o’clock, at 4 o’clock and 8 o’clock and averages across biological replicates. Arteriosclerosis and tubulosclerosis was estimated by averaging the scoring between 5 representative 250µm^2^ areas within each slide. Scoring was performed as follows: 0 – no visible changes, 1 – 50% of area shows changes (e.g. tubular vacuolization, thickening of vessel walls), 2 – >75% area shows changes (e.g. tubular atrophy, vessel atrophy or onion shapes).

#### Albuminuria Assay Quantification

Images were processed using FiJi (v1.54). Autofluorescence is shown in gray scale. DAPI-channel was not shown. Images were processed into maximum-intensity projections after background subtraction (rolling ball radius 50 pixel). After pseudo-flat field correction albumin-positive area was thresholded using “default” method and measured. Total kidney area was measured using Autofluorescence. Albumin positive area was calculated relative to total kidney area. A total of 10 young male fish and 7 old fish were used per age group for age-dependent quantification. For experimental quantification 4 males per group and 3 females per condition were analyzed.

#### Kidney Tubule Ex vivo Measurements

Fluorescence intensities were quantified from images using FiJi (version 1.54). Briefly, two or three adjacent cellular and luminal areas are selected from each tubule and the pixel intensity for each area was calculated. The mean intensity value was averaged between multiple tubules of the same condition. Averaged mean fluorescence for cells and lumen were divided and cell/lumen ratio was calculated to interpolate tubule transport.

#### CD31 positive Area Quantification

Images were analyzed using FiJi (v 1.54). First, a Maximum Intensity Projection was generated. Then a pseudo-flat field correction was performed (Gaussian Blur = 125 pixels). “Enhance Local Contrast (CLAHE)” and a Gaussian Blur of 2 pixels were applied to all images. Total kidney area was calculated by measuring area after maximum intensity thresholding using the default method. Masks for CD31 positive area were generated using “Analyze particles” with a size restriction of “0.1 microns – infinity”. Total CD31 positive Area was divided by total kidney area for each biological replicate. A total of 7 young males, 4 old males, 3 young females and 9 old females were used.

#### 3D Analysis of Vessel Networks

Image Analysis was adapted from Bonda et al., 2020. In summary, we performed a background subtraction (rolling ball radius =50µm) and selected 3 random 250µm^2^ square areas per kidney as representatives for unbiased data analysis. Amira software (version 2024.2) was used to generate a median mask (using 3D and 10 iterations) with a gamma correction of 2. Images were binarized with Fiji using the Otsu thresholding method on auto mode. Back in Amira, the *multi-thresholding* module was used to label white pixels as vessels. Small objects were removed using the *Remove Islands* function in 3D with “multiple” close neighbors enabled. Subsequent steps included *Chamfer distance mapping* (interpretation set to XY planes) and vessel skeletonization using *Thinner* module (length of ends set to 5, 10 iterations). Thinned images were converted using *Trace Lines* module into a spatial graph, which was smoothed using *Smooth Line Set* module. Thickness of segments was computed using the *Eval on Lines* command. The output was saved as .mv3D file and imported into WinFiber3D to visualize the spatial graph and to export all vessel network, vessel length, diameter, volume and branch points information for each vessel.

### Statistical data analysis

Data were analyzed in GraphPad Prism (version 10.4.2.). Outlier removal was performed (Q=1%). Normality was assessed via Shapiro-Wilk tests. Normally distributed data were analyzed using unpaired t-tests or one-way ANOVA with Tukey’s post hoc test. Non-parametric data were analyzed using Mann-Whitney or Kruskal-Wallis tests. Each data point represents an individual fish with at least three independent measurements. Data are presented as mean ± standard deviation, with significance thresholds of *p* ≤ 0.05 (**), p ≤ 0.01 (**), and p ≤ 0.001 (****). Graphs displayed were generated in RStudio (v 4.4.3).

### Single nuclei RNA isolation

#### Isolation protocol

We established and verified our own single nuclei RNA isolation protocol for killifish kidneys. Our protocol was established by combining two existing protocols, one for Killifish brain (Teefy et al., 2022) and a kidney-specific protocol for mice (Leiz et al., 2021). For the experiment the following numbers of fish were used: young female n=9, young males n=6, old females n=6, old males n=5. Pooling of multiple kidneys was necessary to achieve an average tissue weight of 20 mg. Young time points were defined as 6 weeks old animals; old time point was defined by the median lifespan of the cohort (in this case: 20 weeks of age).

All old fish were transcardially perfused with ice-cold PBS before kidney collection ^2^. Kidneys were pooled into RNA-later solution and stored at 4°C for 24h. Then all remaining liquid was removed, and tissue was moved to -80°C until isolation.

Nuclei isolation: On day of experimentation tissue was removed from the cryotube, rehydrated with ice-cold PBS on ice. PBS was removed and tissue was resuspended in Nuclei Lysis Buffer (NLB: 10mM Tris-HCl (pH 7.4), 10mM, NaCl, 3mM MgCl_2_, 0.01875% v/v NP-40, 0.2U/µl RNase Inhibitor). Tissue in buffer was transferred into a 2ml dounce homogenizer on ice and homogenized with pestle A for 30 sec. The homogenate was passed through 70µM strainer into a 15 ml tube, previously precoated with 5% BSA overnight. The filter was washed with another 1ml of NLB. The dissociated tissue was moved back into the douncer and homogenized using pestle B for 1 minute. The homogenate was then strained through a 40µm strainer into a 15 tube precoated with 5% BSA. The homogenate was diluted with Nuclei wash buffer (NWB: 2% BSA in PBS, 0.2U/µl RNase Inhibitor) up to 5 ml and gently inverted. The suspension was centrifuged at 500g at 4°C for 10 minutes.

Nuclei purification: Nuclei were purified with an additional wash step. The supernatant was discarded by gently tipping over the tube and the pellet was resuspended in 1 ml NWB using a wide bore P1000. The volume was filled up to 5 ml with NWB and the tube centrifuged 500g at 4°C for 5 minutes.

Debris removal: The supernatant was carefully discarded by gently tipping over the tube. The pellet was resuspended in 1 ml NWB with a wide bore P1000. 300µl debris removal solution (Miltenyi Biotec®, USA) was added and slowly mixed. The solution was overlayed with 1ml NWB and centrifuged at 3000g for 10 minutes at 4°C. The top two phases were gently removed. The remaining phase was diluted to 8ml and centrifuged at 1000g for 10 minutes at 4°C. The supernatant was removed and the pellet resuspended in 1 ml NWB. The solution was filtered through a 20µm filter into a 1.5ml low-bind DNA tube.

Samples were processed which have shown an RNA Integrity number of >7 as measured on a Agilent Bioanalyzer using a Bioanalyzer High Sensitivity RNA Analysis Kit (Agilent®, USA).

Nuclei were counted using Acridine Orange/Propidium Iodide Assay on a CellDrop®( DeNovix®, USA) automated cell counter set to “Nuclei AO/PI”.

A Chromium Next GEM Single Cell 3’ Reagent Kit v3.1 (10x genomics) was used for library construction following the user’s manual. Nuclei were concentrated to 1000nuclei/µl for a targeted cell recovery of 7000.

#### Single Nuclei Sequencing Analysis

The transcriptomic data supporting the findings of this study are openly available in the Gene Expression Omnibus (GEO) under accession number GSE297623 (<https://www.ncbi.nlm.nih.gov/geo/query/acc.cgi?acc=GSE297623>). *During peer review the files are accessible using a token, which will be given upon request by the corresponding author.*

#### Data Processing

The raw fastq-format sequencing data was processed using nf-core/scrnaseq (v2.6.0) (doi: <https://doi.org/10.5281/zenodo.3568187>) of the nf-core collection of workflows (doi: <https://doi.org/10.1038/s41587-020-0439-x>) built on the Nextflow framework (doi: <https://doi.org/10.1038/nbt.3820>). The reads were aligned to the Ensembl (v111) African turquoise killifish reference using STAR (doi: <https://doi.org/10.1093/bioinformatics/bts635>) implemented in the 10x Genomics Cell Ranger software.

#### Data Analysis with Seurat

R Studio was used to run custom R scripts for analysis. Genes expressed in fewer than three cells were excluded from the analysis. Cell quality control was conducted on the data by eliminating the bottom 10% of cells based on read count and gene count, along with cells containing a mitochondrial percentage greater than 10%. Doublets were identified and removed using the *DoubletFinder* package (v2.0.4). (doi: https://doi.org/10.1016/j.cels.2019.03.003). The *Seurat* package (v5.0.2) (doi: <https://doi.org/10.1016/j.cell.2021.04.048>) was used for clustering and cell type identification. Expression levels were normalized through the “NormalizeData” function with the LogNormalize method. For visualization, the “ScaleData” function was used to regress out differences in the number of molecules, number of genes, percent mitochondrial genes, and cell cycle effects. Principal component analysis (PCA) was conducted, and the first 43 PCs were used to identify clusters and generate both t-distributed stochastic neighbor embedding (t-SNE) and uniform manifold approximation and projection (UMAP) plots.

#### Cell type identification

UMAPs were displayed in Loupe Browser 8 (10x Chromium), and specific cell clusters were determined by the expression of marker genes that define specific cell types and known expression as known in zebrafish, mice and humans. Identified cell cluster were verified by using FindAllMarkers function with options min.pct = 0.25, min.diff.pct=0.25.

#### Data Analysis with Cellchat

The *CellChat* package (v2.1.2) (doi: <https://doi.org/10.1038/s41596-024-01045-4>) was used to infer intercellular communication within the single-nuclei data. The pre-curated human database for intercellular communication was translated into African turquoise killifish using orthology maps derived from Ensembl (v111). Non-one-to-one mappings were maintained in the translated database by duplicating interaction entries for each African turquoise killifish gene that was mapped to a Human gene. This was done to eliminate the chance of not detecting existing interactions.

With the African turquoise killifish *CellChat* database tested and intact, normalized counts were extracted, and each condition was independently analyzed for intercellular communication. Comparisons were then made by merging individual runs of *CellChat.*
