## Supplementary figures and images for "SGLT2 Inhibition Ameliorates Age-Dependent Renovascular Rarefaction"

### Supplementary Figure S1

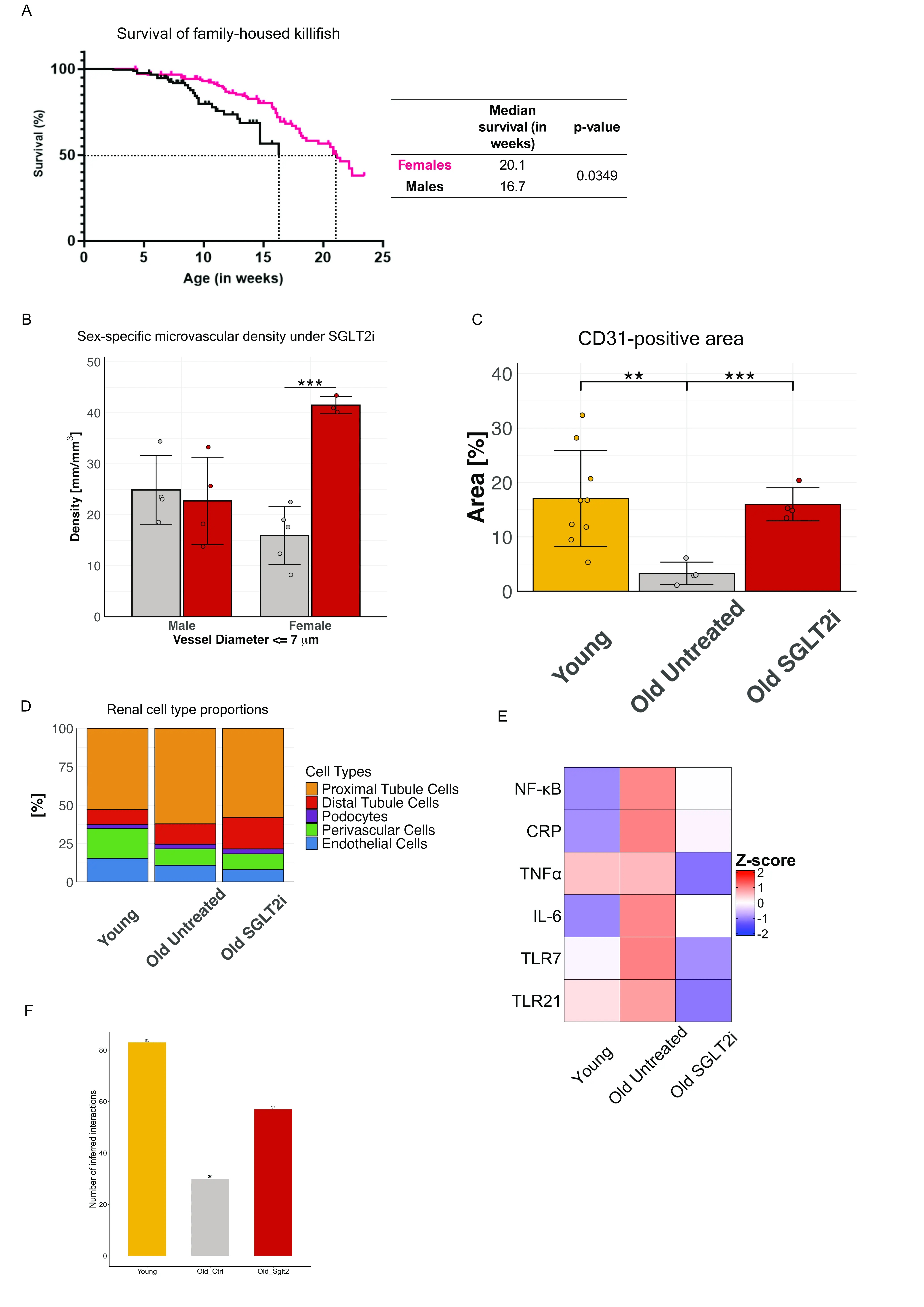

### Supplementary Figure S2

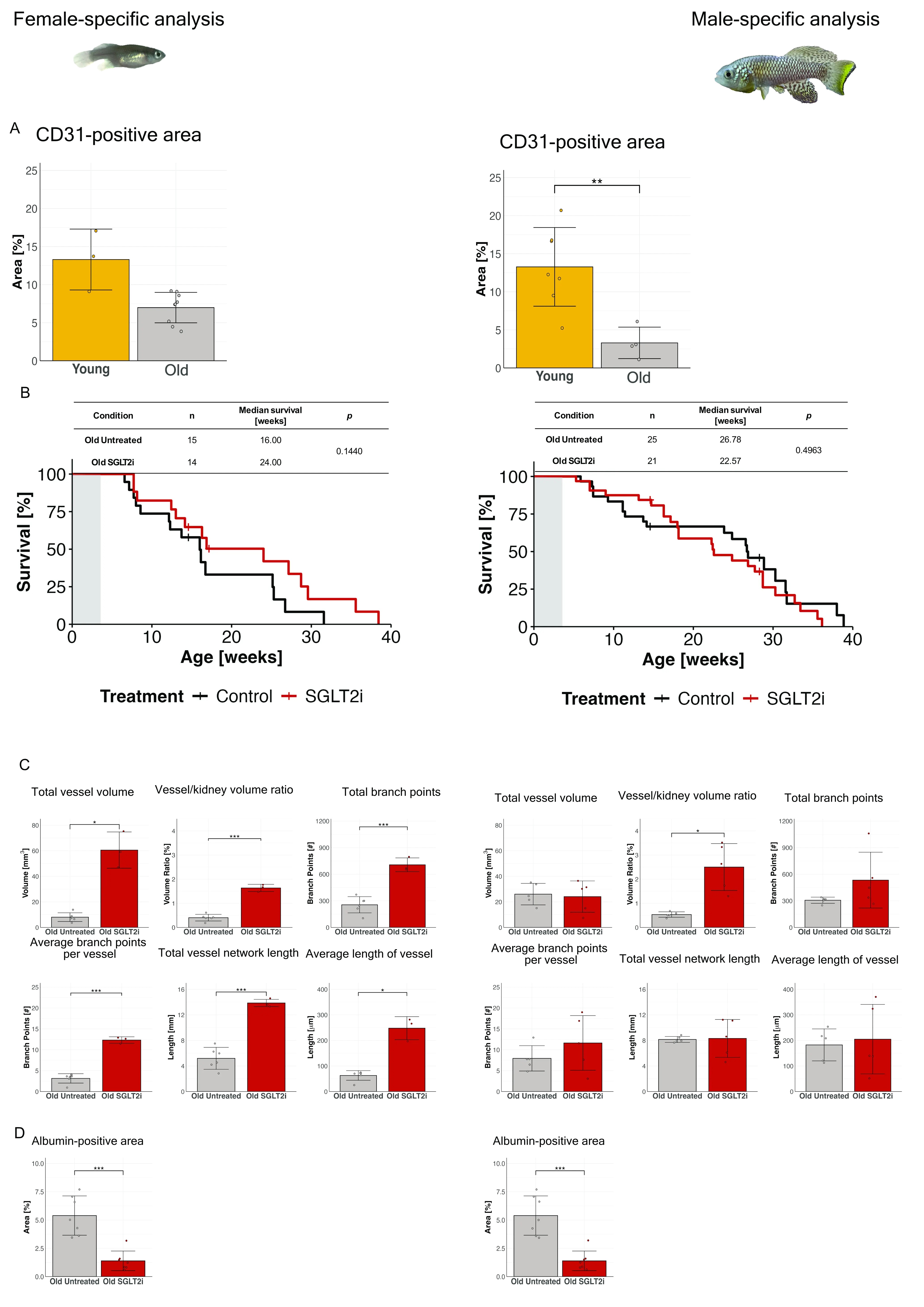

### Supplementary Figure S3

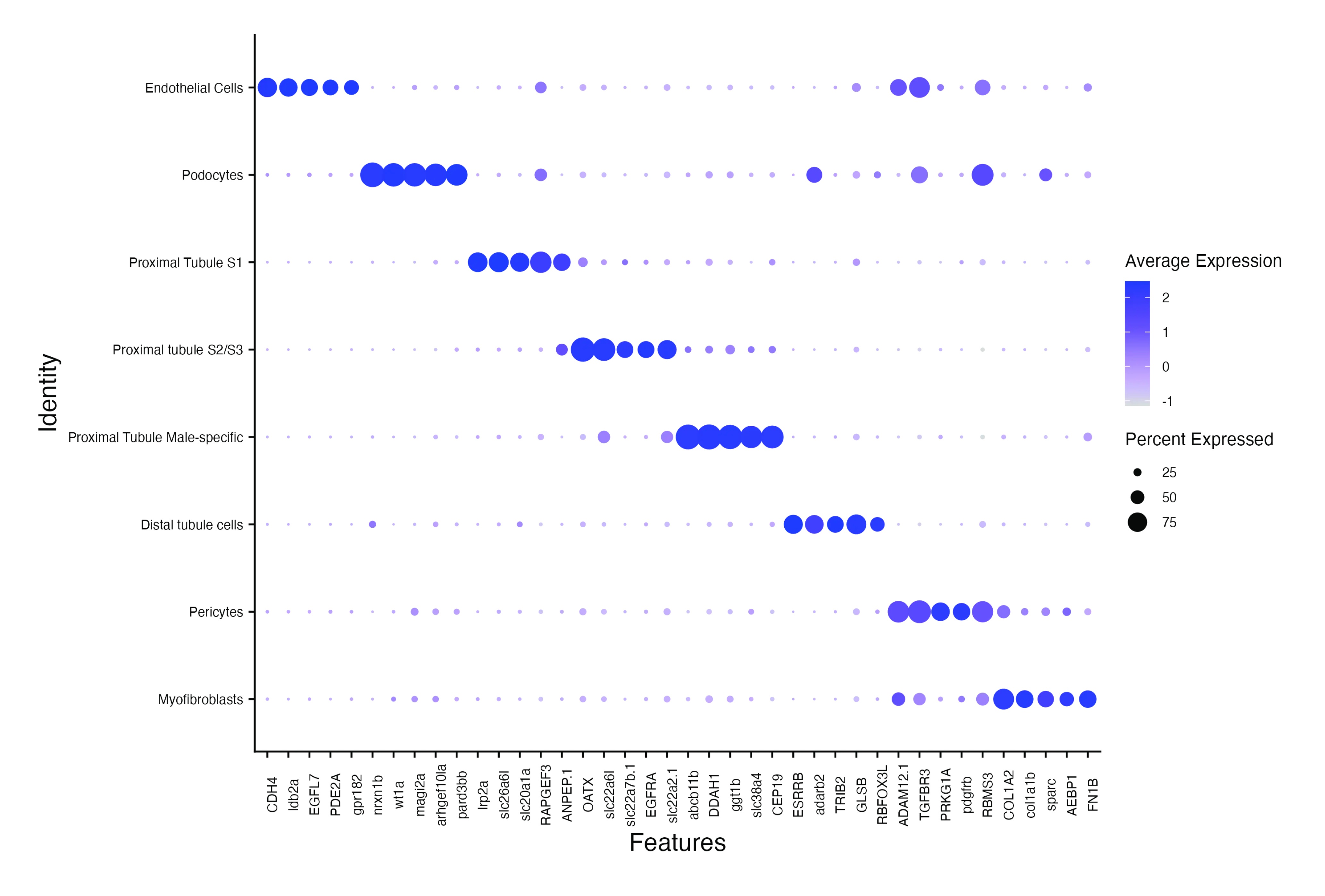

### Supplementary Figure S4

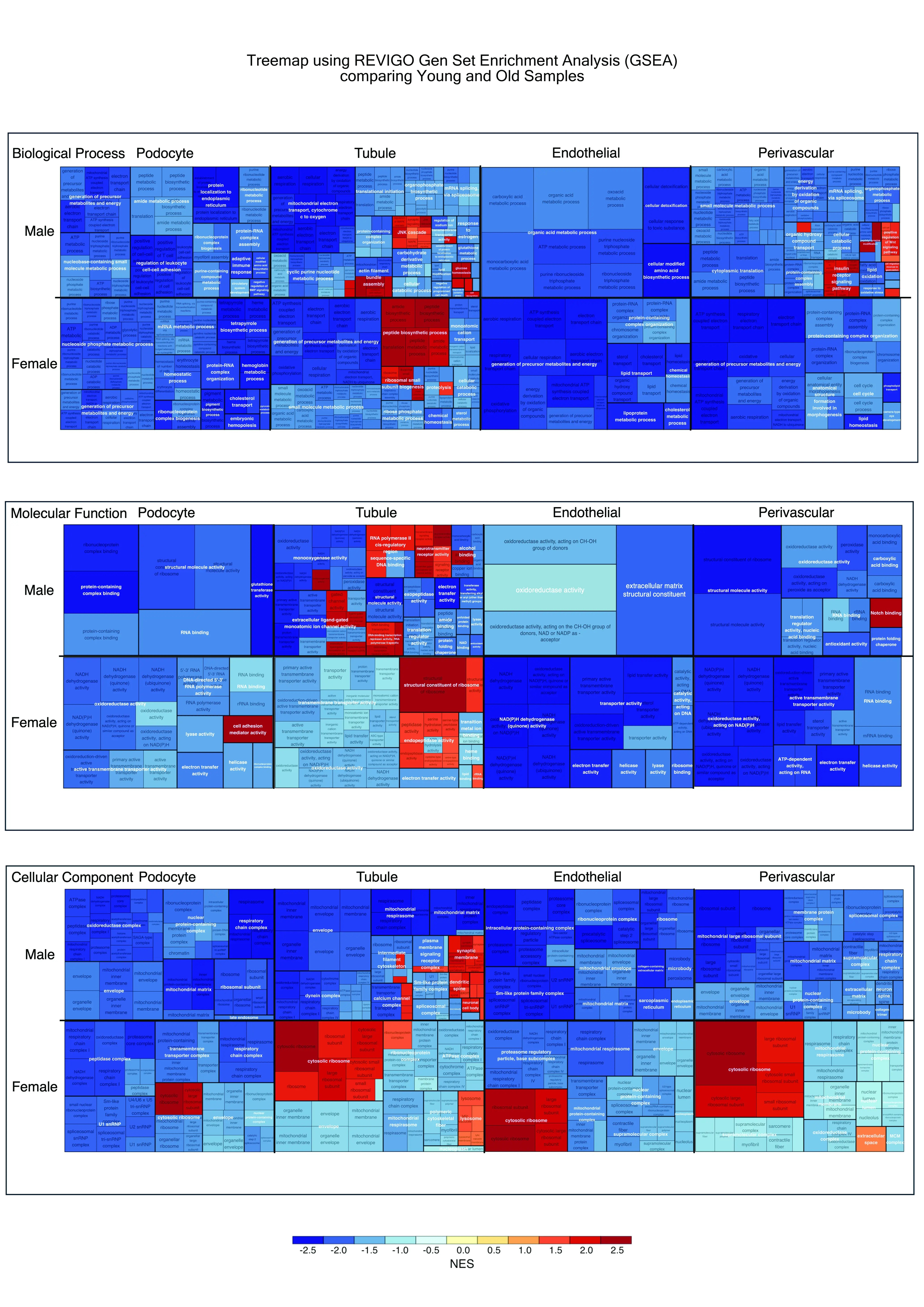

### Supplementary Figure S5

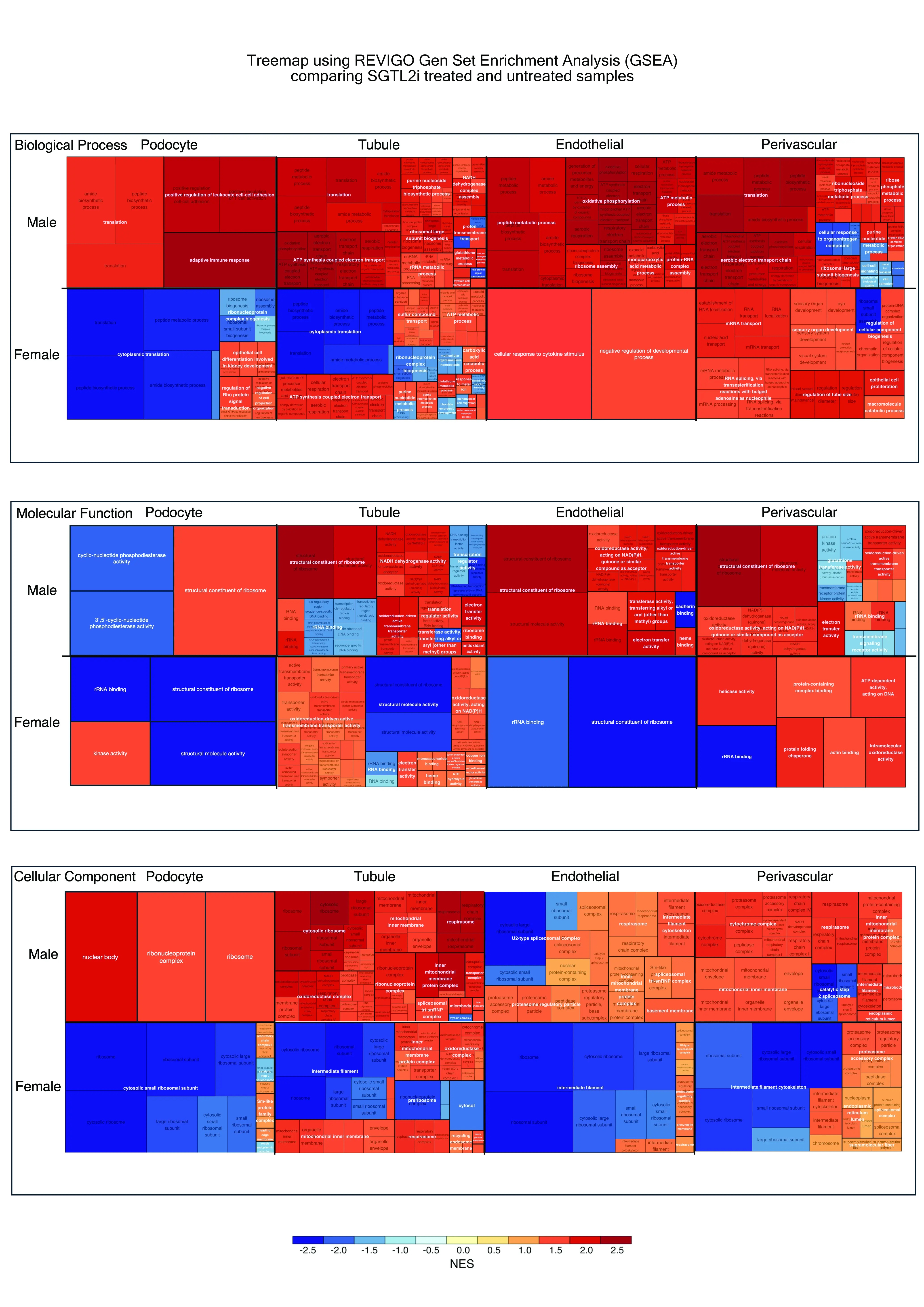
